# Characterization of Outer Membrane β-barrel Proteins of *Pasteurella multocida* for Multi-Epitope Vaccine Design against Human Pasteurellosis

**DOI:** 10.64898/2026.05.28.728361

**Authors:** Amisha Panda, Jahnvi Kapoor, Sanjiv Kumar, Anannya Bandyopadhyay

## Abstract

*Pasteurella multocida* is a facultative anaerobic, Gram-negative coccobacillus that causes pasteurellosis in companion animals, livestock, and poultry and poses a significant zoonotic risk to humans through bite wounds, scratches, licking, and transfer of bodily fluids. Although vaccines are available for livestock and poultry, no vaccine is currently licensed for human use. In this study, we systematically identified and characterized 29 outer membrane β-barrel (OMBB) proteins in *P. multocida* Past9 proteome and classified them into functional categories, including TonB-dependent receptors, porins, autotransporters, adhesins, and efflux pumps. B-cell, cytotoxic T-lymphocyte (CTL), and helper T-lymphocyte (HTL) epitopes were predicted from the identified proteins and screened based on antigenicity, non-allergenicity, and non-toxicity. Epitopes conserved across eight human-infecting *P. multocida* strains and located within the extracellular loop (ECL) region were incorporated into a multi-epitope vaccine (MEV) construct. The designed MEV was predicted to be antigenic, non-allergenic, and soluble. Its tertiary structural model was iteratively refined and validated. Molecular docking with human toll-like receptors 4/2 (TLR4/TLR2) predicted stable interactions, further supported by 100 ns molecular dynamics simulations. Immune simulation of the MEV construct predicted a strong simulated immune response. Furthermore, codon optimization and in silico cloning supported the feasibility of recombinant MEV expression in *E. coli*. The construct was benchmarked against OmpH, a well-known antigenic protein and exhibited broadly comparable predicted immune responses and receptor-binding energetics. This study proposes a designed MEV candidate against human pasteurellosis and highlights OMBB proteins as potential immunogenic targets for vaccine development.

**Graphical Abstract:** 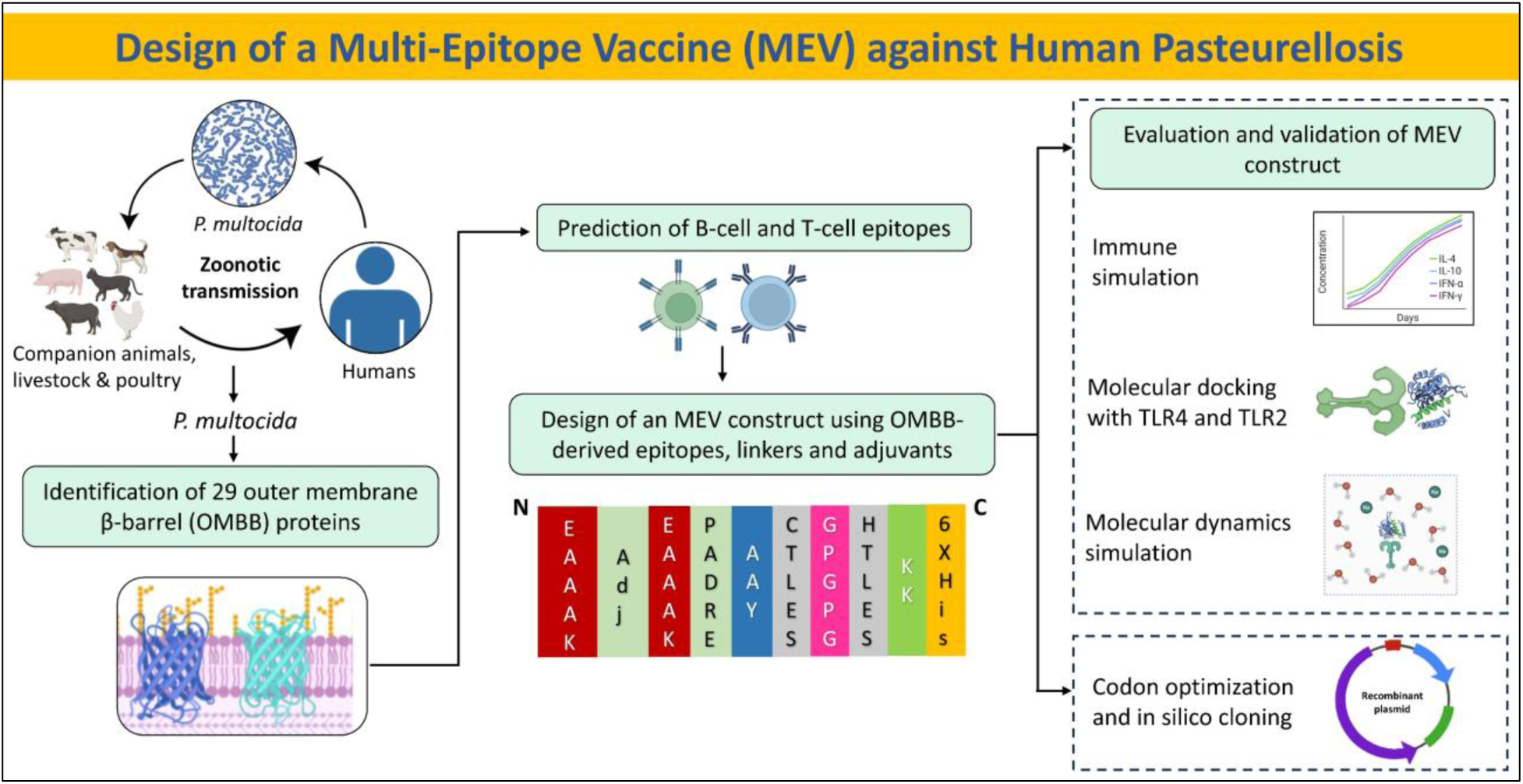

## 1. Introduction

Pasteurellosis caused by the bacterium *Pasteurella multocida* (Pm) infects a wide range of animals, including companion animals (cats and dogs), livestock and poultry, leading to substantial economic losses ^1^. Pm is an opportunistic, facultative anaerobic pathogen that commonly resides in the nasopharynx of animals. It is most commonly transmitted to humans through bite wounds, scratches, licking and transfer of bodily fluids ^2^. The bacterium is found in 50% of dog bites and 75% of cat bites ^3^. The disease affects ∼20-50% of the 1-2 million individuals bitten annually in the United States ^4^ and is endemic in tropical regions like South Asia, Southeast Asia, and Africa ^5^. Pm can also be transmitted from an infected mother to her newborn ^6^. Pm infections are characterized by skin and soft tissue infections, endocarditis, septicemia, meningitis, urinary tract infections, and respiratory infections ^3^. Current treatment relies primarily on antibiotic (penicillin) therapy; however, Pm multidrug-resistant strains are increasingly being reported ^2,7,8^. Therefore, vaccination remains the most affordable and promising strategy to combat pasteurellosis. Current vaccines, widely used in clinical practices for livestock and poultry, include inactivated and attenuated vaccines ^9^; however, no vaccine is currently available for humans, underscoring the need to develop a vaccine candidate against human pasteurellosis.

Pm is a Gram-negative, non-motile coccobacillus belonging to the family Pasteurellaceae ^2^. It possesses a typical diderm cell envelope comprising an outer membrane (OM), inner membrane (IM), and periplasmic space ^10^. The OM consists of an inner phospholipid layer and an outer lipopolysaccharide (LPS) leaflet ^11^. The bacterium is serologically classified into five common serogroups (A, B, D, E, and F) based on the composition of its polysaccharide capsule, and into 16 LPS serovars according to Carter’s and Heddleston’s classification methods ^12–15^. It has a single circular chromosome of ∼2.4 Mb that encodes 2241 proteins. Like other Gram-negative bacteria, Pm possesses a diverse repertoire of outer membrane β-barrel (OMBB) proteins ^16,17^. β-barrels are cylindrical proteins of various sizes comprising 8 to 36 anti-parallel β-strands ^18^. OMBB proteins mediate host-pathogen interactions in addition to their essential roles in nutrient uptake, membrane biogenesis, and OM assembly ^19^. Earlier computational work identified 98 OMPs in Pm avian strain Pm70 and 107 OMPs in porcine strain 3480 ^20^. Another proteomic level study of non-human Pm isolates identified 22 OMPs, including porins, adhesins, and iron-uptake receptors ^21^. Surface-exposed OMBB proteins are highly accessible to the host immune system, which makes them ideal vaccine candidates. Several OMP-based subunit vaccines targeting PlpE, OmpH, Omp87, and VacJ have conferred 66.7-100% protection against Pm serotypes A and B in mice ^22–25^. Moreover, a multi-epitope vaccine (MEV) proposed for swine pasteurellosis using six Pm proteins, including three OMBB proteins (PlpE, OmpA, OmpH, VacJ, Omp87, and Cp39), demonstrated protective immune responses against multiple Pm serotypes in mice ^26^. Recently, an in silico designed MEV based on PlpE, OmpA, OmpH, and LolA was proposed for ovine pasteurellosis ^27^. These findings support the potential of surface-exposed OMBB proteins as promising candidates for MEV development against human pasteurellosis.

In the present study, we predicted OMBB proteins in Pm and designed an MEV construct using the predicted proteins. We employed a consensus-based computational framework that comprises nine different prediction tools to identify OMBB proteins ^28,29^. Structural models were generated using five modeling tools and aligned to assess concordance. The identified OMBB proteins were classified into functional categories, and their amino acid sequence variations across eight human-infecting Pm strains were analyzed. Immunogenic potential of the identified proteins was assessed by predicting B-cell, cytotoxic T-lymphocyte (CTL), and helper T-lymphocyte (HTL) epitopes. An MEV construct was designed using surface-exposed B-cell, CTL, and HTL epitopes, which were linked with appropriate linkers and supplemented with adjuvants ^30^. The tertiary structural model of MEV was docked with human toll-like receptors 4 and 2 (TLR4/2) to assess binding interactions, followed by molecular dynamics (MD) simulations to confirm the stability of the complexes. Immune simulation was performed to assess the potential of the MEV to elicit an immune response. Furthermore, to enhance recombinant protein expression in *E. coli*, codon optimization and in silico cloning were performed. This study highlights the potential of OMBB proteins as candidates for developing an MEV against Pm infection, offering a promising strategy to tackle this pathogen.

## 2. Materials and Methods

### 2.1 Prediction of OMBB proteins

Pm subsp. *septica* Past9 strain, a well-characterized human-infecting strain, was selected for this study ^31^. We downloaded the amino acid sequences of all 2241 proteins of Pm Past9 proteome from NCBI (http://www.ncbi.nlm.nih.gov/accessed) using the genome assembly ASM3522456v1 (accessed on 4 July 2025) ^32^.

The computational pipeline employed in this study was adapted from a previously established framework ^30^. A detailed description of the methodology is given below and is schematically shown in Figure 1. Nine computational tools were used to analyze the complete proteome file of Pm (Table S1). Pepstats tool (https://www.ebi.ac.uk/jdispatcher/seqstats/emboss_pepstats) (accessed on 7 July 2025) of the EMBOSS package was used to determine the peptide length, molecular weight, charge, and isoelectric point (pI) for all protein sequences ^33^. SPAAN (https://sourceforge.net/projects/adhesin/files/SPAAN/spaan_64_bit.tar.gz/download) (accessed on 8 July 2025) was employed to predict the adhesins or adhesin-like proteins ^34^. SignalP 5.0, based on a deep convolutional and recurrent neural network architecture (https://services.healthtech.dtu.dk/services/SignalP-5.0/) (accessed on 14 July 2025), was used to predict signal peptide and its cleavage position in the protein sequences ^35^. CELLO v.2.5 (http://cello.life.nctu.edu.tw/) (accessed on 24 July 2025) ^36^ and PSORTb v3.0 (https://www.psort.org/psortb/) (accessed on 24 July 2025) ^37^ were used to determine the subcellular localization of the proteins.

**Fig. 1.**
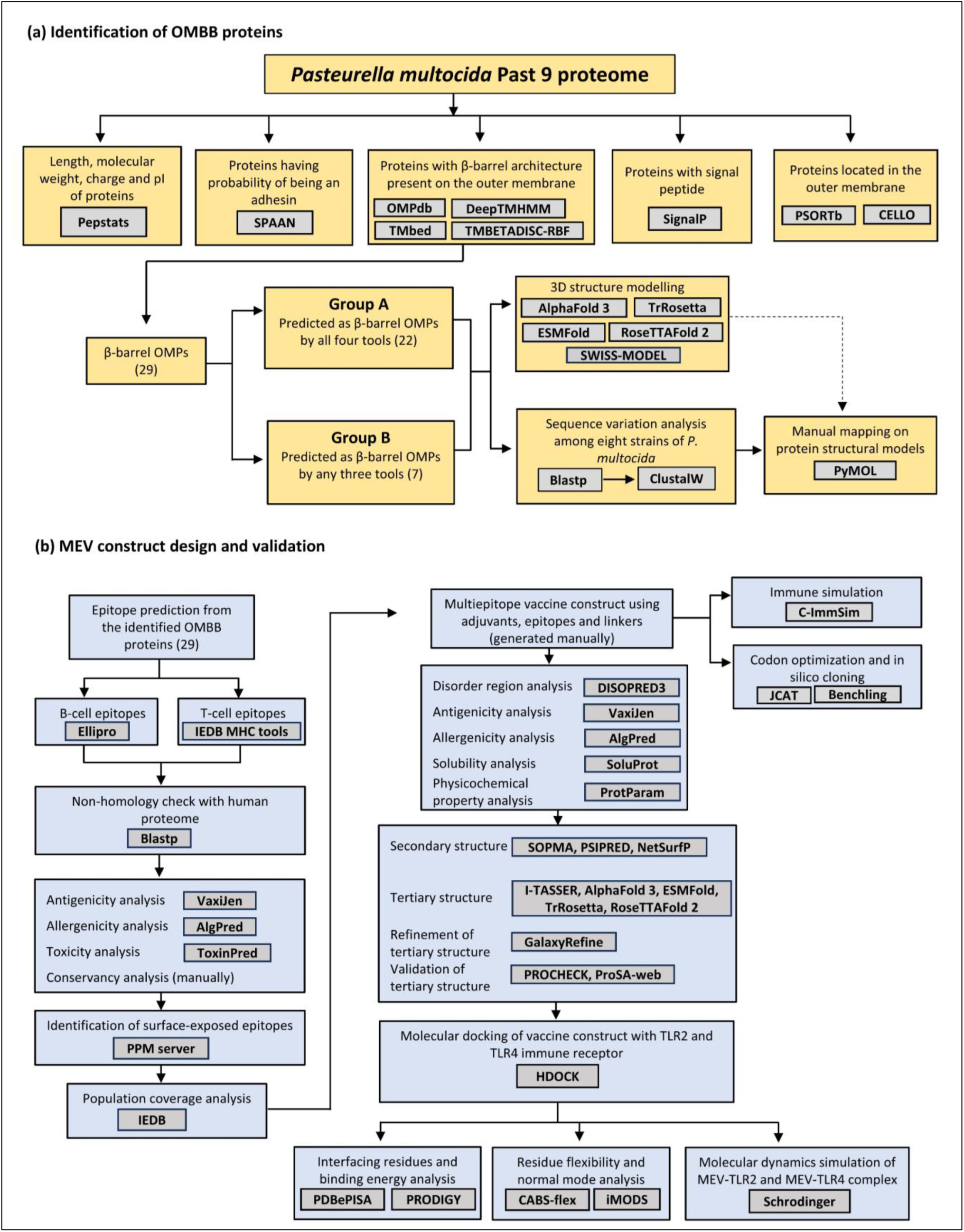
Computational workflow for outer membrane β-barrel (OMBB) proteins identification and multi-epitope vaccine design. **(a)** Framework for identification of OMBB proteins. **(b)** Workflow employed for designing a multi-epitope vaccine against human pasteurellosis.

A consensus-based computational framework was used to identify OMBB proteins, where the outputs from four OM localization and β-barrel prediction tools: one database-OMPdb (http://aias.biol.uoa.gr/OMPdb/) (accessed on 24 July 2025) ^38^, and three tools-DeepTMHMM (https://dtu.biolib.com/DeepTMHMM) (accessed on 24 July 2025) ^39^, TMBETADISC-RBF (http://rbf.bioinfo.tw/∼sachen/OMP.html) (accessed on 27 July 2025) ^40^, and TMbed (https://github.com/BernhoferM/TMbed) (accessed on 27 July 2025) ^41^ was taken into consideration for identification of OMBB proteins. Furthermore, proteins were categorized based on the number of tools providing positive outputs for a given protein sequence. This implied that higher confidence was given to those predicted as OMBB protein by a greater number of tools.

### 2.2 Structural modeling and validation of predicted OMBB proteins

Structural models of the predicted proteins in Group A and Group B were generated using AlphaFold 3 server (https://alphafoldserver.com/) (accessed on 11 December, 2025) ^42^. To validate the structural models, we generated 3D structural models using four additional modeling tools: ESMFold (https://colab.research.google.com/github/sokrypton/ColabFold/blob/main/ESMFold.ipynb) (accessed on 12 December, 2025) ^43^, SWISS-MODEL (https://swissmodel.expasy.org/) (accessed on 2 January, 2026) ^44^, RoseTTAFold (https://robetta.bakerlab.org/) (accessed on 19 January, 2026) ^45^ and TrRosetta (https://yanglab.qd.sdu.edu.cn/trRosetta/) (accessed on 20 January, 2026) ^46^. For each protein, the top-ranked model, based on the highest confidence score, was selected for visualization and further analysis. The resulting atomic coordinate files were visualized using PyMOL ^47^. Structural models were aligned using US-align (https://zhanggroup.org/US-align/) (accessed on 22 January, 2026) to assess structural similarities and differences across models ^48,49^. US-align is a universal structure alignment tool designed to compare and align 3D macromolecular structures. Alignments were visualized using PyMOL, and corresponding root mean square deviation (RMSD) values were noted (Table S2).

### 2.3 Amino acid sequence variation analysis across Pm strains

Predicted OMBB protein sequences from Pm Past9 were compared across eight complete Pm proteomes to assess sequence variations (Table S3). Eight Pm human-infecting strains were considered for amino acid sequence variation analysis. Orthologous sequences for each protein were identified using BLASTP (E-value < 1.0E-03, bitscore > 100) (https://blast.ncbi.nlm.nih.gov/Blast.cgi?PROGRAM=blastp&PAGE_TYPE=BlastSearch&LINK_LOC=blasthome) (accessed on 24 January 2026), followed by Multiple Sequence Alignment (MSA) using Clustal Omega ^50^. MSA analysis identified amino acid variations among the orthologs, which were subsequently mapped onto structural models using PyMOL.

### 2.4 B-cell and T-cell epitopes prediction

ElliPro (http://tools.iedb.org/ellipro/), a webserver of the Immune Epitope Database (IEDB) (accessed on 27 January 2026), was used to predict both linear and conformational B-cell epitopes ^51^. The IEDB webserver predicts B-cell and T-cell epitopes using structure- and sequence-based algorithms trained on experimentally validated epitope data ^52^. Predictions of B-cell epitopes were performed using default parameters. Epitopes longer than 50 amino acids were excluded as they may introduce non-immunogenic flanking regions.

CTL and HTL epitopes were predicted using IEDB MHC I (https://tools.iedb.org/mhci/) and MHC II (http://tools.iedb.org/mhcii/) prediction tools, respectively, using NetMHCpan-4.1 for class I and NetMHCIIpan-4.0 for class II (accessed on 4 February 2026) ^53^. CTL epitopes (9-mer peptides) were predicted with a percentile rank threshold of < 0.5. The analysis included the following Human Leukocyte Antigen (HLA) alleles: HLA-A*01:01, HLA-A*02:01, HLA-A*11:01, HLA-A*24:02, HLA-B*07:02, HLA-B*08:01, HLA-B*15:01, HLA-B*44:02, HLA-C*04:01, and HLA-C*07:02. HTL epitope prediction targeted 15-mer peptides with a percentile rank threshold < 2 for the following HLA alleles: HLA-DRB1*01:01, HLA-DRB1*03:01, HLA-DRB1*04:01, HLA-DRB1*07:01, HLA-DRB1*08:02, HLA-DRB1*13:01, HLA-DRB1*13:02, and HLA-DRB1*15:01.

### 2.5 Homology of predicted epitopes with human proteome

Proteins that are homologous to human sequences are more likely to induce an autoimmune response. Given the fact, a comparison was made between *Homo sapiens* proteome (TaxID: 9606) and predicted peptides using the NCBI BLASTP database (https://blast.ncbi.nlm.nih.gov/Blast.cgi?PROGRAM=blastp&PAGE_TYPE=BlastSearch&LINK_LOC=blasthome) (accessed on 6 February, 2026)^54^. Peptides with E-values exceeding 0.05 were considered non-homologous and selected as potential vaccine candidates. Since BLASTP statistics are less reliable for short peptides, k-mer analysis was performed using PepMatch (https://nextgen-tools.iedb.org/pipeline?tool=pepmatch) (accessed on 31 July 2026) ^55^ as an additional validation step. Peptides exhibiting 100% identity (zero mismatch) to sequences in the human proteome were removed.

### 2.6 Antigenicity, allergenicity and toxicity evaluation of predicted epitopes

The non-homologous epitopes were screened for antigenicity, allergenicity and toxicity. Antigenicity was analyzed using the VaxiJen2.0 online web server (https://www.ddg-pharmfac.net/vaxijen/VaxiJen/VaxiJen.html) (accessed on 1 August 2026) that utilizes the bacterial prediction model and a threshold score of 0.4 ^56^. Allergenicity of the predicted epitopes was assessed using AlgPred 2.0 (https://webs.iiitd.edu.in/raghava/algpred2/) (accessed on 1 August 2026), with default parameters ^57^. Toxicity of the selected epitopes was predicted using the ToxinPred server (http://crdd.osdd.net/raghava/toxinpred/) (accessed on 1 August 2026). Epitopes identified as antigenic, non-allergenic, and non-toxic were selected for further analysis.

### 2.7 Conservation analysis and surface-exposure of predicted epitopes

MSA was performed as described in Section 2.3, and the resulting output was used to evaluate the conservation of selected epitopes across eight strains of Pm. Conserved epitopes across eight Pm strains were selected for further analysis to ensure the broad applicability and effectiveness of the predicted epitopes. Furthermore, PPM 3.0 (https://opm.phar.umich.edu/ppm_server3) (accessed on 5 August 2026) was subsequently employed to validate the membrane orientation of the corresponding OMBB proteins and determine whether the selected epitopes were exposed on the extracellular surface ^58^. Given that a prospective vaccine candidate should be surface-exposed protein that can stimulate the host immune system for clearance of the pathogen, epitopes located on the extracellular loop (ECL) region were considered for further analysis.

### 2.8 Population coverage analysis of T-cell epitopes

The population coverage of the selected CTL and HTL epitopes was estimated using the IEDB Population Coverage tool (https://tools.iedb.org/population/) (accessed on 6 August 2026) based on their corresponding HLA class I and class II alleles ^59^. The analysis included the population dataset of pasteurellosis-endemic regions ^4,5^ to estimate the proportion of the population expected to respond to the selected epitopes.

### 2.9 Development of MEV construct

Antigenic, non-allergenic, non-toxic, conserved, and surface-exposed epitopes were selected for inclusion in the MEV construct. Based on their respective prediction scores/ranks, a total of 12 epitopes (two linear B-cell, five CTL and five HTL epitopes) were selected and joined together using appropriate peptide linkers to construct the vaccine sequence ^60,61^. HTL epitopes were connected using GPGPG linkers, while CTL epitopes were connected with AAY linkers. B-cell epitopes were linked using KK linkers. Two adjuvants-β-defensin 3 (accession no.: Q5U7J2) and PADRE sequence were added at the N-terminus with the help of EAAAK linker to enhance immunogenicity. An additional methionine residue was added at the N-terminus to provide the translation initiation site. A hexa-histidine tag for purification was added at the C-terminus through a KK linker to obtain the complete vaccine protein sequence ^62^.

### 2.10 Evaluation of MEV construct

The MEV was evaluated using four bioinformatics tools-ProtParam (http://web.expasy.org/protparam/) was used to assess physicochemical properties ^63^, VaxiJen 2.0 for antigenicity assessment ^56^, SoluProt (https://loschmidt.chemi.muni.cz/soluprot/) for solubility prediction ^64^ and AlgPred 2.0 for allergenicity evaluation ^57^ (all tools accessed on 9 August 2026). SoluProt employs a machine learning (ML) algorithm to predict the propensity of a protein to be soluble upon expression in *E.coli* ^64^. Intrinsically disordered regions (IDRs) of the MEV construct were predicted using the DISOPRED3 (http://bioinf.cs.ucl.ac.uk/disopred) (accessed on 9 August 2026) that utilizes ML modules combined with support vector machine (SVM)-based classifier ^65^. Previous studies have demonstrated that Pm OmpH is immunogenic and provides protection in different animal models following challenge, highlighting its potential as a vaccine candidate ^66^. Therefore, OmpH of Pm (UniProt Accession no. O54339.1) was used as a positive control.

### 2.11 Structure prediction of the MEV construct and assessment of model quality

Secondary structures of the MEV construct were predicted using the SOPMA (Self-Optimized Prediction method With Alignment) (http://npsa-pbil.ibcp.fr/cgibin/npsa_automat.pl?page=/NPSA/npsa_sopma.html), PSIPRED 4.0 (https://bioinf.cs.ucl.ac.uk/psipred/) and NetSurP 3.0 (https://services.healthtech.dtu.dk/services/NetSurfP-3.0/) (all tools accessed on 9 August 2026). SOPMA employs MSA-based consensus scoring ^67^, PSIPRED utilizes PSI-BLAST-derived evolutionary profiles ^68^, and NetSurfP uses machine-learning models ^69^ to assess the structure of the target protein by predicting the formation of α-helices, β-turns, random coils, and extended strands. Tertiary structure of the MEV was generated using AlphaFold 3 (accessed on 10 August 2026), followed by refinement using GalaxyRefine web server (http://galaxy.seoklab.org/) (accessed on 11 August 2026) to improve model quality ^70^. The quality of the MEV model was assessed using the PROCHECK (https://saves.mbi.ucla.edu/) ^71^ and ProSA-web server (https://prosa.services.came.sbg.ac.at/prosa.php) ^72^ (both tools accessed on 11 August 2026). Model quality was assessed using the Ramachandran plot and Z-score analysis, which provide insights into the overall stability and stereochemical quality of the predicted structures, respectively.

### 2.12 Molecular docking with human TLR4 and TLR2

Molecular docking between the MEV construct and human TLR4/TLR2 was performed to evaluate the interaction with host immune receptors. The atomic coordinate files of human TLR4 (PDB Id: 3FXI) and TLR2 (PDB Id: 6NIG) were obtained from the RCSB PDB database (accessed on 11 August 2026). Docking was performed using the HDOCK server (http://hdock.phys.hust.edu.cn/) (accessed on 11 August 2026). The resulting MEV-TLR4 and MEV-TLR2 complexes were evaluated for interacting amino acid residues using the PDBePISA server (https://www.ebi.ac.uk/pdbe/pisa/) (accessed on 12 August, 2026) ^73^ and the docked complex was visualized using PyMOL. The docked complexes were assessed by calculating interaction energetics using the PRODIGY (PROtein binDIng enerGY prediction) server ^74^ (https://rascar.science.uu.nl/prodigy/) (accessed on 12 August 2026), which provides Gibbs free energy (ΔG, kcal mol^−1^) or dissociation constant (K_d_, M) for intermolecular interactions typically within a 5.5 Å distance cutoff.

### 2.13 Residue flexibility analysis of MEV-TLR4/2 complexes

Flexibility analysis of the docked MEV-TLR4/TLR2 complexes were carried out to evaluate its structural stability, conformational flexibility, and dynamic interactions under near-physiological conditions. CABS-flex v3.0 (https://lcbio.pl/cabsflex3/) (accessed on 12 August 2026) was employed to perform rapid protein flexibility analysis using the coarse-grained CABS model ^75^. The MEV-TLR4/TLR2 coordinate files were submitted to the server for a 10-ns simulation with default settings, generating ten modeled clusters and a root mean square fluctuation (RMSF) profile for per-residue fluctuations. It also provides RMSD values for medoid (representative structure of a cluster) in comparison with the input reference structure. The iMODS server (https://imods.iqf.csic.es/) (accessed on 12 August 2026) was used to perform normal mode analysis (NMA)-based simulation, assuming that the lowest-frequency modes represent the largest, functionally relevant motions ^76^. Docked PDB structures were analyzed with default parameters, and stability was assessed using deformability, B-factor, eigenvalues, variance, covariance map, and elastic network analyses.

### 2.14 MD simulation of MEV-TLR4/2 complexes

MD simulations of the docked MEV-TLR4/2 complexes were performed to assess the structural stability using the Desmond module in the Schrodinger suite 2025-3 (accessed on 18 August 2026). The docked complexes were first prepared using the Protein Preparation Wizard in Schrodinger Suite 2025-3 ^77^, followed by energy minimization using the OPLS4 force field. The system was prepared using the System Builder tool, where each protein-ligand complex was solvated with TIP3P water molecules and enclosed with an orthorhombic simulation box. The OPLS4 force field at physiological pH 7.4 was applied, and Na+ and Cl-ions were added to neutralize the system and maintain a salt concentration of 0.15M. MD simulations were performed under NPT conditions at a bar pressure of 1.01325 and a constant temperature of 300K. The resulting trajectories were analyzed using Simulation Interactions Diagram (SID) to obtain RMSD plot for each protein-ligand complex.

### 2.15 Immune simulation of the MEV construct

The online web server C-ImmSim (https://kraken.iac.rm.cnr.it/C-IMMSIM/) (accessed on 8 September 2026) was used to study the extent and type of immune response generated by the MEV construct in humans ^78^. The server simulates the immune response using PSSMs and machine-learning methods and evaluates interactions between the antigen and various immune cells. At the same time, it simulates three different anatomical regions in mammals: bone marrow, thymus, and lymph nodes ^79^. The vaccine was administered in three doses at three different time steps: 1, 84, and 168, with each time step corresponding to 8 hours. The simulation parameters were set as follows: random seed 12345, simulation volume of 50, and simulation steps of 1050 and HLA alleles selected for the analysis were: HLA-A*01:01, HLA-A*02:01, HLA-B*07:02, HLA-B*08:01, HLA-DRB1*01:01 and HLA-DRB1*03:01.

### 2.16 Codon optimization and in silico cloning

The designed MEV construct was codon optimized to ensure efficient heterologous expression in *E. coli* K-12 using Java Codon Adaptation Tool (JCat) (http://www.jcat.de/) (accessed on 16 August 2026) ^80^. This improves codon usage in accordance with *E. coli* codon preference while maintaining the amino acid sequence of the vaccine construct. To facilitate efficient transcription and translation, rho-independent transcription terminators, prokaryotic ribosome binding sites, and restriction enzyme cleavage sites were avoided. The optimized nucleotide sequence was evaluated for its codon adaptation index (CAI) and GC content.

For in silico cloning, BamHI and EcoRI restriction enzyme recognition sites were introduced at the 5′ and 3’ termini of the codon-optimized nucleotide sequence, respectively. The optimized MEV gene was then inserted into the pET-28a (+) expression vector between the BamHI and EcoRI restriction sites using the molecular cloning tool available in Benchling (https://www.benchling.com/) (accessed on 16 August 2026). The recombinant plasmid construct was analyzed to confirm the correct orientation and in-frame insertion of the MEV sequence for potential recombinant protein expression in *E. coli*.

## 3. Results

### 3.1 Prediction of OMBB proteins

A consensus-based computational framework was employed to identify OMBB proteins in Pm Past9, a well-characterized human-derived isolate ^31^. The designed pipeline utilized nine computational tools: Pepstats, SPAAN, SignalP, CELLO, PSORTb, OMPdb, DeepTMHMM, TMBETADISC-RBF, and TMbed. Predictions from all the tools were consolidated for each protein sequence (Table S1). Outputs from four OMBB prediction tools-OMPdb, DeepTMHMM, TMBETADISC-RBF, and TMbed-were considered for OMMB protein identification. The identified proteins were classified into two groups: Group A constituted proteins predicted as OMBB by all four prediction tools, and Group B comprised those predicted by three of the four tools (Table 1). Consensus prediction initially predicted 23 proteins in Group A and nine in Group B. AlphaFold 3 generated structures were visualized in PyMOL, and proteins having β-barrel architectures were retained. Following manual curation, 29 OMBB proteins were identified, of which 22 belonged to Group A, and seven to Group B. AlphaFold 3-generated models were validated using four additional modeling tools: ESMFold, SWISS-MODEL, RoseTTAFold, and TrRosetta. Structural similarity between the generated models was assessed using the US-align server, which yielded RMSD values ranging from 1.27 to 5.28 Å (Table S2).

**Table 1:** Identified OMBB proteins using consensus-based computational approach.

| Protein categories* | Protein accession no. | Protein annotation (NCBI)<sup>‡</sup> |
| --- | --- | --- |
| **Group A (22 proteins)** |  |  |
| TonB-dependent receptors | WP_096743251.1 | TonB-dependent receptor domain-containing protein |
|  | WP_104228050.1 | TonB-dependent receptor plug domain-containing protein |
|  | WP_126417535.1,<br>WP_249984729.1,<br>WP_249984736.1 | TonB-dependent hemoglobin/transferrin/lactoferrin family receptor |
| Porins | WP_046333314.1 | Porin OmpA |
|  | WP_249984777.1 | Opacity family porin |
|  | WP_324023089.1,<br>WP_324023090.1 | Porin |
| OM assembly proteins | WP_083004621.1 | Outer membrane protein assembly factor BamA |
|  | WP_126417754.1 | LPS assembly protein LptD |
|  | WP_249984417.1 | Surface lipoprotein assembly modifier |
|  | WP_324022736.1 | Autotransporter assembly complex family protein |
| Autotransporters | WP_249984510.1 | Autotransporter domain-containing protein |
|  | WP_249984830.1 | Autotransporter outer membrane beta-barrel domain-containing protein |
| Enzymatic OMPs | WP_115299510.1 | Sialidase family protein |
|  | WP_324022762.1 | Phospholipase A |
| Secretion system OMPs | WP_083004763.1,<br>WP_324023075.1 | ShlB/FhaC/HecB family hemolysin secretion/activation protein |
| Adhesin | WP_249984768.1 | MipA/OmpV family protein |
| Transporter | WP_108511463.1 | Outer membrane protein transport protein |
| OM β-barrel protein | WP_005757734.1 | Outer membrane beta-barrel protein |
| **Group B (7 proteins)** |  |  |
| TonB-dependent receptors | WP_249984435.1,<br>WP_249984789.1 | TonB-dependent receptor |
|  | WP_324023022.1 | TonB-dependent receptor domain-containing protein |
| Efflux pump OMPs | WP_005756417.1 | Toxin/drug exporter Tde |
|  | WP_104227660.1 | Efflux transporter outer membrane subunit |
| Porin | WP_083005347.1 | OmpW family outer membrane protein |
| Enzymatic OMP | WP_249984615.1 | Sialidase family protein |

<sup>‡</sup> Proteins were named as per their annotation in the NCBI database, searched using protein accession number.
\*Outputs from four OMBB-specific prediction tools- OMPdb, DeepTMHMM, TMBETADISC-RBF, and TMbed- were taken into consideration for the classification of OMBB proteins (Table 1). Group A included proteins predicted as OMBB by all four prediction tools, and Group B included proteins predicted as OMBB by three of the four tools.

The identified proteins were classified into ten functional categories based on their NCBI annotations-TonB-dependent receptors, porins, OM assembly proteins, autotransporters, enzymatic OMPs, secretion system OMPs, adhesins, transporters, efflux pump OMPs and OM β-barrel proteins (Table 1).

#### 3.1.1 TonB-dependent receptors

This group included eight proteins: WP_096743251.1, WP_104228050.1, WP_126417535.1, WP_249984729.1, WP_249984736.1, WP_249984435.1, WP_249984789.1, and WP_324023022.1. TonB-dependent proteins such as HgbA and TbpA have been characterized in Pm as mediators of iron acquisition. These proteins facilitate heme uptake from haemoglobin and iron transport across the OM, respectively ^81,82^. Structural models for these eight proteins, generated using five tools, revealed 22-stranded β-barrel architecture containing a plug domain at the N-terminal that undergoes conformational changes to regulate the opening and closing of the pore (Figure 2). In Pm, TonB-dependent receptor proteins are involved in the transport of various molecules, such as iron, haemoglobin, transferrin, lactoferrin, and vitamin B12 ^83^.

**Fig. 2.**
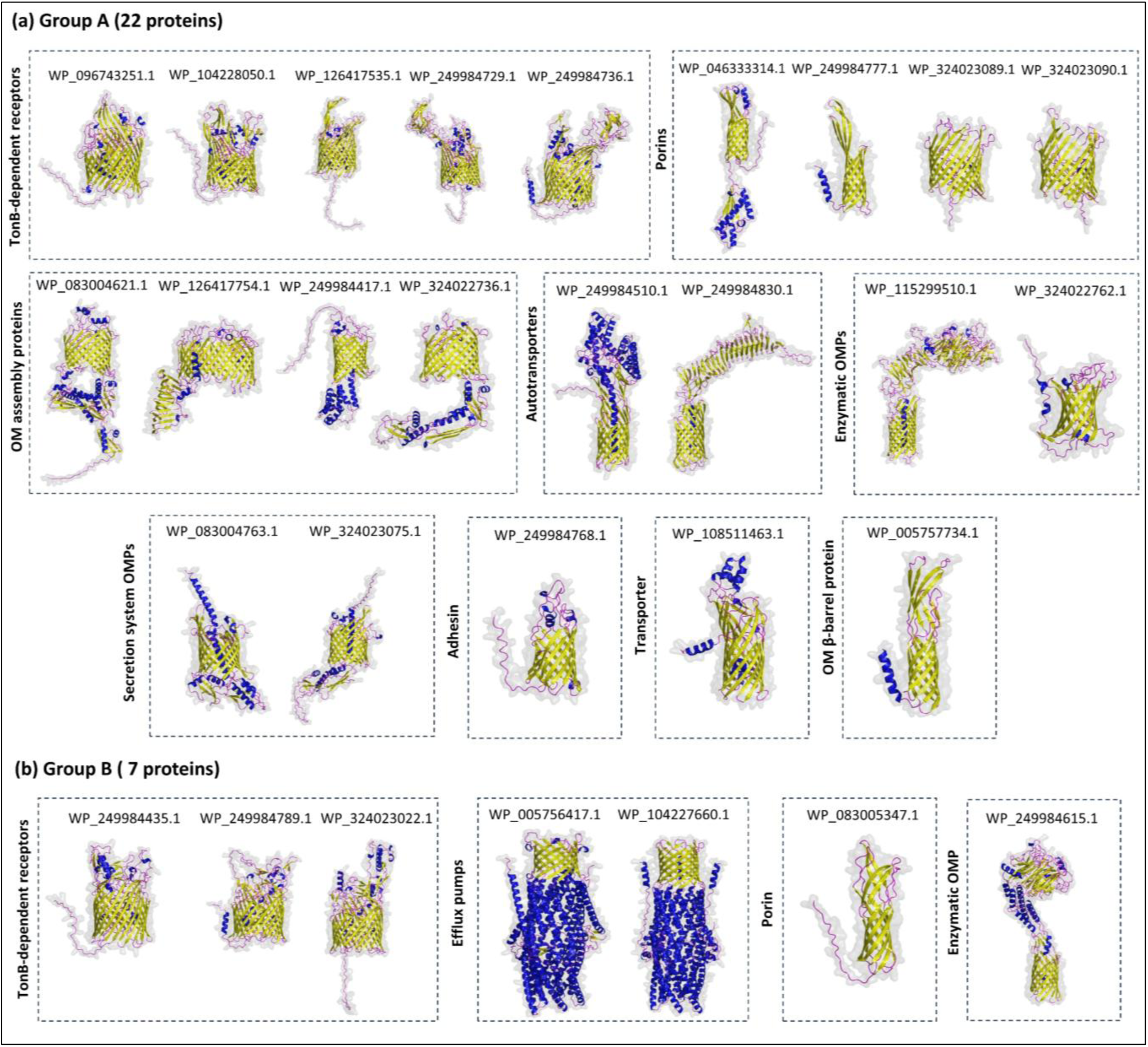
Structural models of β-barrel outer membrane proteins. **(a)** Structural models of 22 Group-A OMBB proteins generated using AlphaFold 3. **(b)** AlphaFold 3-generated models of seven Group-B OMBB proteins.

#### 3.1.2 Porins

Five proteins annotated as porin family proteins are-porin OmpA (WP_046333314.1), opacity family porin (WP_249984777.1), two general porins (WP_324023089.1 and WP_324023090.1) and OmpW family protein (WP_083005347.1) (Table 1). Porin OmpA mediates the interaction of Pm with host cells by binding to heparin and fibronectin, thereby functioning as a crucial invasion factor ^84,85^. The generated OmpA model exhibited an eight-stranded β-barrel structure (Figure 2a), consistent with the previously reported structure of Pm OmpA ^86^. Opacity (Opa) proteins are a polymorphic family of ∼28 kDa OMPs that act as adhesins and invasins in *Neisseria gonorrhoeae* ^87,88^, and its predicted model showed an eight-stranded β-barrel structure (Figure 2a). In contrast, general porins were predicted to form 16-stranded β-barrel architecture (Figure 2a). Pm OmpW family protein is known to have an eight-stranded β-barrel architecture, and our study validated this prediction by generating models from five prediction tools. Pm OmpW forms a long and narrow hydrophobic channel for the transport of hydrophobic molecules and is highly conserved across Pm strains ^89,90^.

#### 3.1.3 OM assembly proteins

This class comprised four proteins-WP_083004621.1, WP_126417754.1, WP_249984417.1, and WP_324022736.1. WP_083004621.1 is annotated as outer membrane protein assembly factor BamA. The structural model generated for this protein showed a 16-stranded β-barrel architecture and five periplasmic polypeptide transport-associated (POTRA) domains (Figure 2a). The predicted β-barrel domain also contained a lateral gate between strands β1 and β16. Overall, the structure closely resembles the typical BamA protein found in Gram-negative bacteria ^91^. WP_126417754.1 is a lipopolysaccharide (LPS) transport protein D (LptD). Our structural model predicted a 26-stranded β-barrel structure with a distinctive periplasmic β-jelly roll domain (Figure 2a), highly similar to the crystallized LptD structures of *Shigella flexneri* and *N. gonorrhoeae* ^92,93^. WP_249984417.1, annotated as a surface lipoprotein assembly modifier, was predicted to form a 14-stranded β-barrel architecture with an estimated molecular weight of 58.3 kDa (Figure 2a). WP_324022736.1 is annotated as an autotransporter assembly complex family protein and was predicted to form a 16-stranded β-barrel with three POTRA domains (Figure 2a), which resembles Translocation and Assembly Module subunit A (TamA) protein found in Gram-negative bacteria ^94^. All four proteins are currently annotated only in NCBI and have not been experimentally characterized.

#### 3.1.4 Autotransporters

The autotransporter class included two proteins: WP_249984510.1 and WP_249984830.1, both predicted to possess an N-terminal extracellular passenger domain and a C-terminal 12-stranded β-barrel domain (Figure 2a). Although structural evidence for these proteins in Pm has not been reported, their predicted architectures are similar to those of autotransporters in other Gram-negative bacteria. The extracellular passenger domains of autotransporters are known to be antigenic ^95^, highlighting these proteins as promising targets for vaccine development.

#### 3.1.5 Enzymatic OMPs

This group contained three proteins: WP_115299510.1, WP_324022762.1, and WP_249984615.1. WP_115299510.1 and WP_249984615.1 are annotated as sialidase family proteins, which are crucial virulence factors that hydrolyze sialic acids from host mucosal surfaces, thereby facilitating bacterial colonization and infection ^96^. Predicted structural models for these two proteins showed 12-stranded β-barrel architecture comprising an N-terminal extracellular passenger domain and a C-terminal β-barrel domain (Figure 2). In contrast, WP_324022762.1, annotated as phospholipase A, was predicted to form a 12-stranded β-barrel structure without an extracellular passenger domain (Figure 2a). This protein is annotated only in NCBI and lacks experimental characterization.

#### 3.1.6 Secretion system OMPs

This class included two proteins: WP_083004763.1 and WP_324023075.1, which are annotated as ShlB/FhaC/HecB family hemolysin secretion/activation proteins. In Gram-negative bacteria, ShlB/FhaC/HecB family proteins are OM two-partner secretion (TPS; Type Vb) transporters that help secrete and activate hemolysin or adhesin partner proteins (ShlA, FhaB, and HecA) ^97,98^. Both proteins were predicted to adopt 16-stranded β-barrel structure (Figure 2a) with an N-terminal α-helix traversing the transmembrane barrel and two POTRA domains, which closely resembles the reported *Bordetella pertussis* FhaC structure (PDB ID: 3NJT) ^99^.

#### 3.1.7 Adhesin

WP_249984768.1 is annotated as MipA/OmpV family protein. Our study predicted a 12-stranded β-barrel architecture (Figure 2a). OmpV family protein helps in adhesion and invasion of *Salmonella typhimurium* to intestinal epithelial cells and thus plays a vital role in pathogenesis ^100^. This protein is annotated only in NCBI and remains experimentally uncharacterized in Pm.

#### 3.1.8 Transporter

WP_108511463.1, annotated as “outer membrane protein transport protein”, was predicted to have a 14-stranded β-barrel structure (Figure 2a). In Pm, the protein has been annotated only in NCBI, with no experimental characterization reported.

#### 3.1.9 OM β-barrel protein

WP_005757734.1, annotated as an outer membrane β-barrel protein, is estimated to have a molecular weight of 26.5 kDa. The structural model predicted an eight-stranded β-barrel structure (Figure 2a).

#### 3.1.10 Efflux pump OMPs

Two proteins: WP_005756417.1 and WP_104227660.1 belong to efflux pump OMPs. These two Pm TolC homologues have been shown to confer resistance to a range of chemicals, including rifampin (512-fold) and acridine orange (128-fold) ^101^. Both were predicted to form trimeric 12-stranded α/β-barrel (Figure 2b).

### 3.2 Amino acid sequence variation analysis

Sequence variation analysis was performed to assess conservation and divergence in proteins, which may reflect host adaptation, host specificity, and virulence. This was performed for the identified OMBB proteins across eight human-infecting Pm strains (Table S3). These variations were identified and mapped onto the corresponding structural models. Amino acid sequence variations were observed to be distributed throughout the proteins, including the extracellular loop (ECL), transmembrane (TM), and intracellular loop (ICL) regions. Out of 29 OMBB proteins, 12 proteins (WP_104228050.1, WP_126417535.1, WP_249984729.1, WP_249984736.1, WP_046333314.1, WP_324023090.1, WP_249984830.1, WP_115299510.1, WP_324023075.1, WP_249984435.1, WP_249984789.1, and WP_249984615.1) have more than 100 variations (Table S4).

### 3.3 Prediction and evaluation of B-cell epitopes

ElliPro was used to predict both linear and conformational B-cell epitopes from all 29 OMBB proteins. Initially, 377 linear B-cell epitopes (LBEs) with a score above 0.5 were predicted and epitopes longer than 50 amino acids were discarded. The remaining LBEs were then screened for homology with human proteome that retained only 151 non-homologous epitopes, of which 33 epitopes were antigenic, non-allergenic and non-toxic. Eight epitopes were conserved across eight Pm strains and mapped onto their corresponding protein models embedded in bacterial OM using PPM 3.0 server to assess surface-exposure. Among them, two LBEs were located in the ECL region and were therefore selected for inclusion in the MEV design (Table 2, Figure 3). Furthermore, 178 conformational B-cell epitopes (CBE) were predicted, of which 20 are listed in Table S5 based on the highest score predicted by the tool. Since the antigenicity of discontinuous CBEs depends on their native protein fold, which is difficult to maintain in a synthetic vaccine construct, only LBEs were considered for MEV design ^102^.

**Fig. 3.**
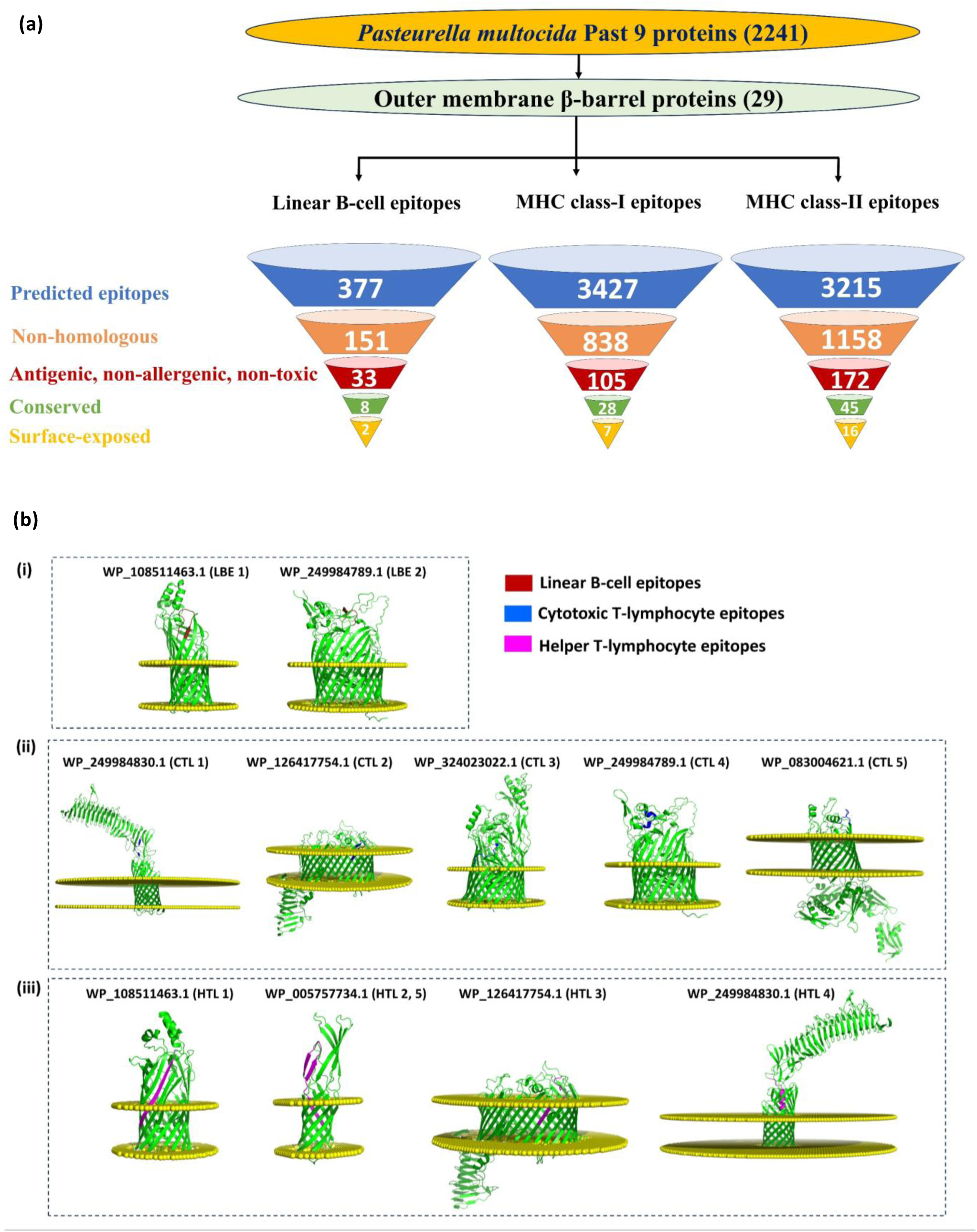
Screening and selection of the predicted epitopes for incorporation into the MEV design. **(a)** Sequential filtering of the predicted epitopes based on non-homology, antigenicity, allergenicity, toxicity, sequence conservation across human-infecting strains, and surface exposure. **(b)** Two linear B-cell, five MHC class-I, and five MHC class-II epitopes were selected based on their surface-exposure and ranking scores, which were mapped onto corresponding protein models. The yellow dummy atoms delineate the outer membrane, where the upper and lower boundaries correspond to the extracellular and periplasmic sides, respectively.

**Table 2:**
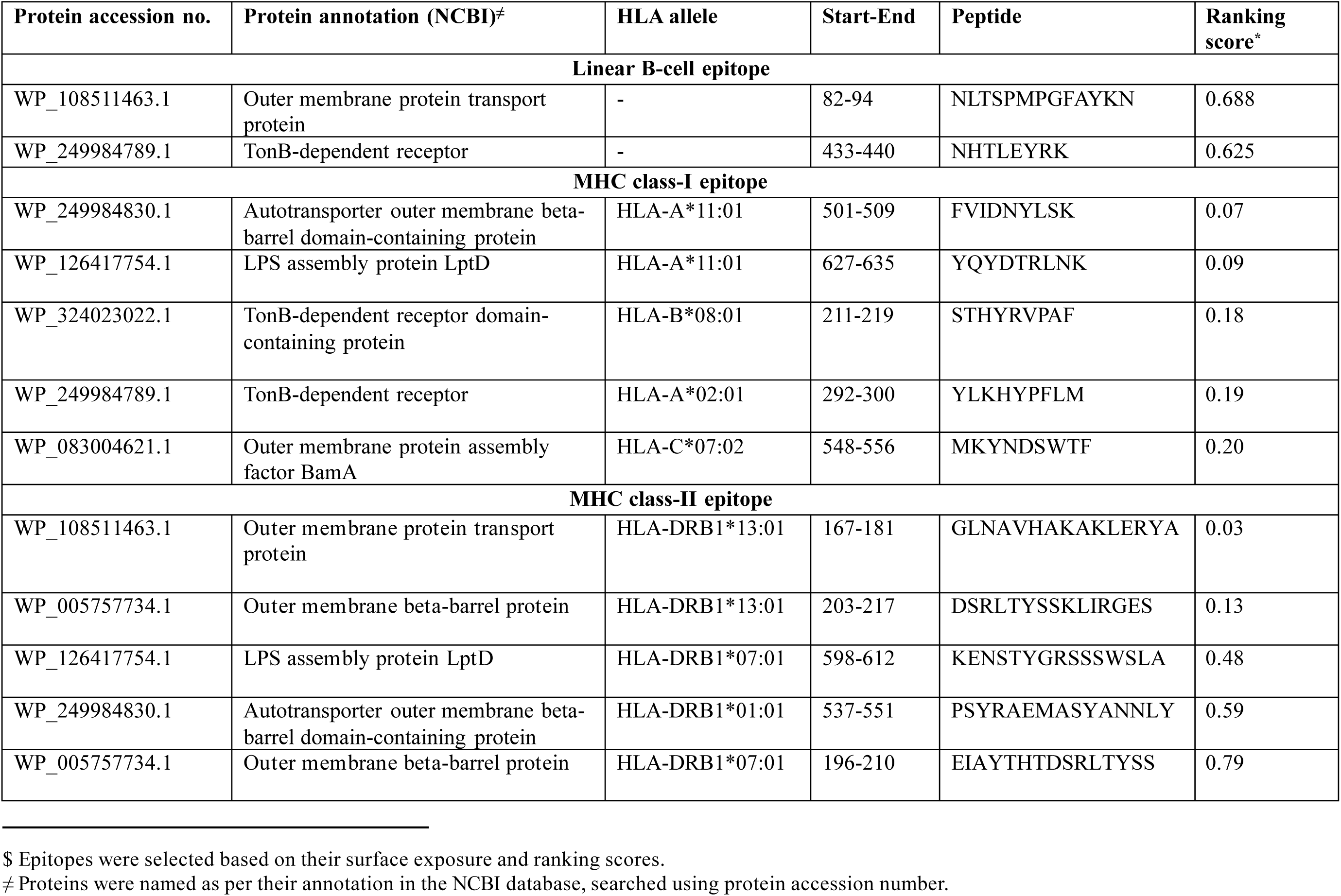
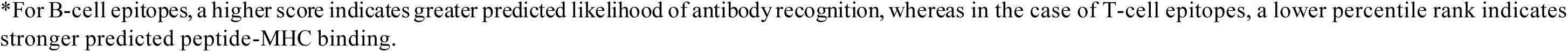
Selected linear B-cell, MHC class-I and MHC class-II epitopes for MEV design^$^.

### 3.4 Prediction and evaluation of T-cell epitopes

Using the IEDB MHC class-I and MHC class-II prediction tools, a total of 3427 CTL epitopes (9-mer) and 3215 HTL epitopes (15-mer) were predicted, respectively. Among these, 28 CTL and 45 HTL epitopes were identified as antigenic, non-allergenic, non-toxic, non-homologous to human proteome and conserved across eight Pm strains. Of these, 7 CTL and 16 HTL epitopes were located within the ECL regions. The five epitopes, each of CTL and HTL, based on NetMHCpan percentile rank, were selected for incorporation into the MEV design (Table 2, Figure 3). None of the identified amino acid variations were present within the epitopes (B-cell, CTL and HTL) selected for MEV design.

The selected CTL and HTL epitopes demonstrated broad population coverage across the evaluated populations, ranging from 79.43% in East Africa to 98.38% in the United States (Figure 4). The estimated global population coverage of selected T-cell epitopes was 97.56%, which suggested that selected epitopes may provide broad HLA-mediated immune recognition across diverse populations (Figure 4).

**Fig. 4.**
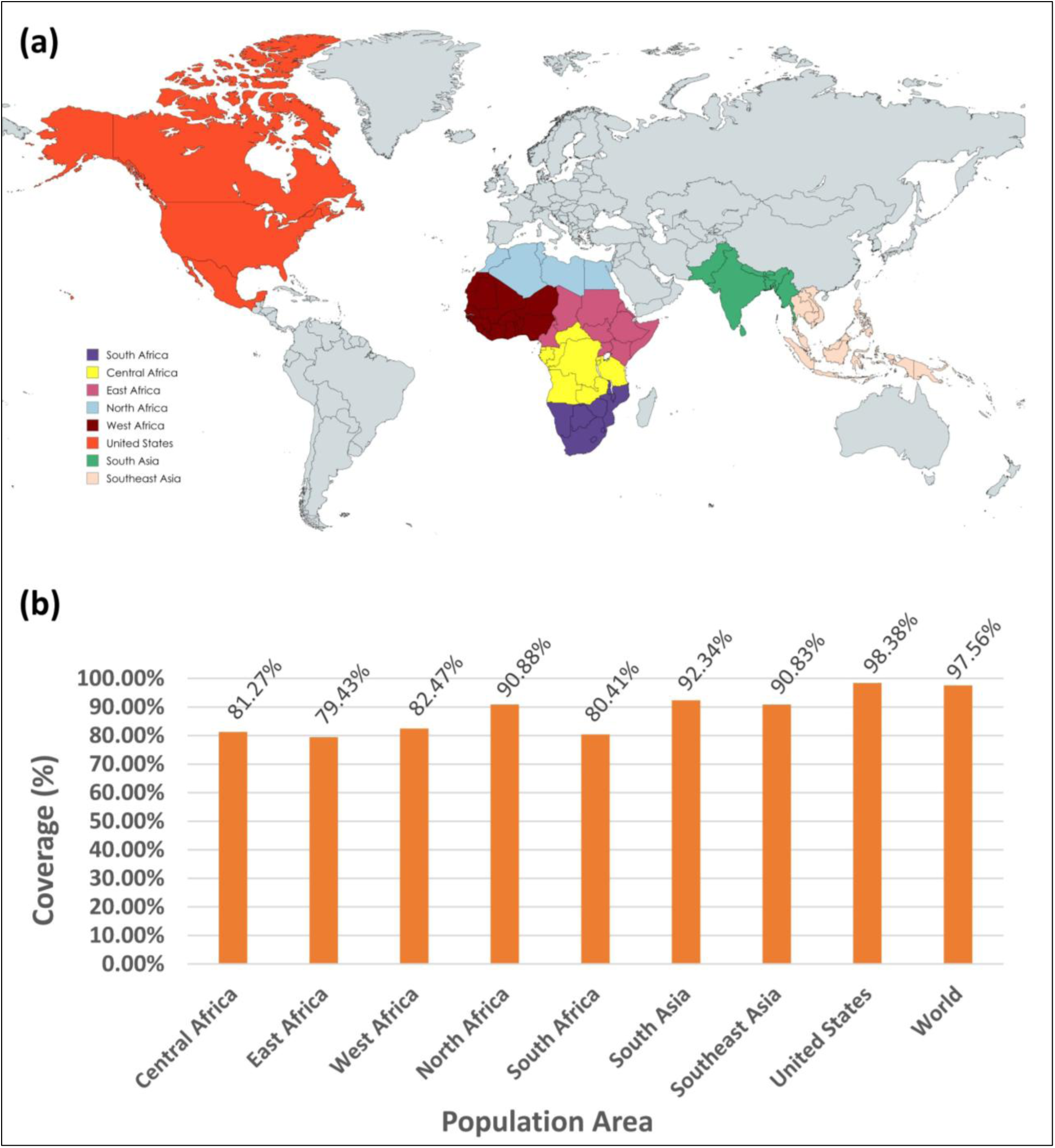
Population coverage analysis of the selected epitopes for MEV design. **(a)** Geographic distribution of population groups included in the analysis. **(b)** Predicted population coverage of the selected epitopes across the pasteurellosis-endemic regions and global population.

### 3.5 Design of the MEV construct and evaluation of physicochemical properties

The screened epitopes (two LBE, five CTL and five HTL) derived from seven OMBB proteins were assembled into a 257 amino acid long MEV construct using flexible (GPGPG, AAY, and KK) and rigid (EAAAK) linkers (Figures 5a and S1). Flexible linkers contain Gly and Ser residues that enhance structural flexibility and promote favorable interdomain interactions, whereas rigid linkers maintain spacing and minimize undesirable interactions. The MEV construct was supplemented with an additional methionine residue and two adjuvants-β-defensin 3 and PADRE peptides. β-defensin 3 acts as an antimicrobial agent by inducing pro-inflammatory cytokine responses from dendritic cells ^103^. PADRE acts as a universal T-helper epitope that can induce robust, broad-spectrum MHC class-II responses and Th1 polarization ^104^. The MEV construct is estimated to have a molecular weight of 28.619 kDa (Table 3), which is below the suggested molecular weight of 110 kDa for MEV constructs ^105^. The theoretical isoelectric point (pI) of the vaccine was calculated to be 9.86, which indicated that it is basic. It is acceptable, as most recombinant protein-based vaccines have pI values ranging from 5 to 10 ^106,107^. The instability index (II), calculated using ProtParam, was 38.26 (threshold < 40), which suggested that the MEV is a stable protein. The hydrophilic nature of the protein was assessed based on the grand average of hydropathicity (GRAVY) score of −0.712. In addition, MEV showed an antigenicity score of 0.8581 (threshold > 0.4) and was found to be non-allergenic. SoluProt predicted a solubility score of 0.831 (threshold > 0.5), suggesting that the protein will remain soluble upon expression in *E. coli* (Table 3). Additionally, residues 244-257 of the MEV were predicted to form IDR as sequences were derived from flexible and surface-exposed epitopes of the parent OMBB proteins (Figure S2). OmpH, which is known to be immunogenic and provide protection in mice against Pm^66^, was used as a positive control in our study. OmpH was also predicted to be stable, hydrophilic, and non-allergenic, with antigenicity comparable to that of the MEV construct (0.7201) but lower solubility (0.535). OmpH exhibited a lower instability index (12.98 vs 38.26) and a higher aliphatic index (78.41 vs 53.74) than the MEV (Table 3), which is attributable to the engineered amino acid composition of the MEV containing multiple epitopes, linkers, and an adjuvant sequence.

**Fig. 5.**
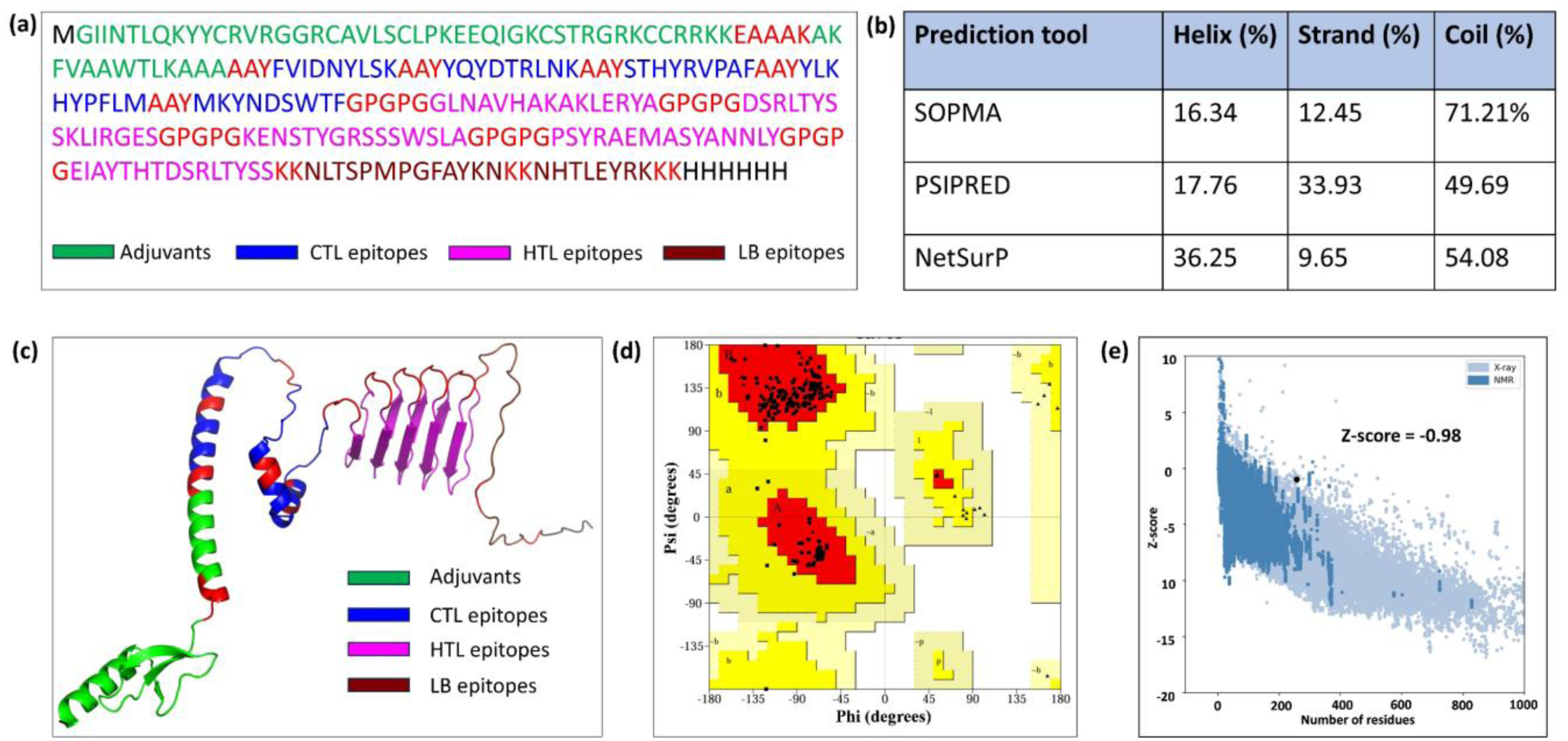
Evaluation of the MEV construct. **(a)** Schematic representation of the MEV construct amino acid sequence. **(b)** Predicted secondary structures of the MEV construct. **(c)** Refined tertiary structural model of the MEV construct. **(d)** Ramachandran plot analysis. **(e)** Z-score plot from ProSA-webserver.

**Table 3:**
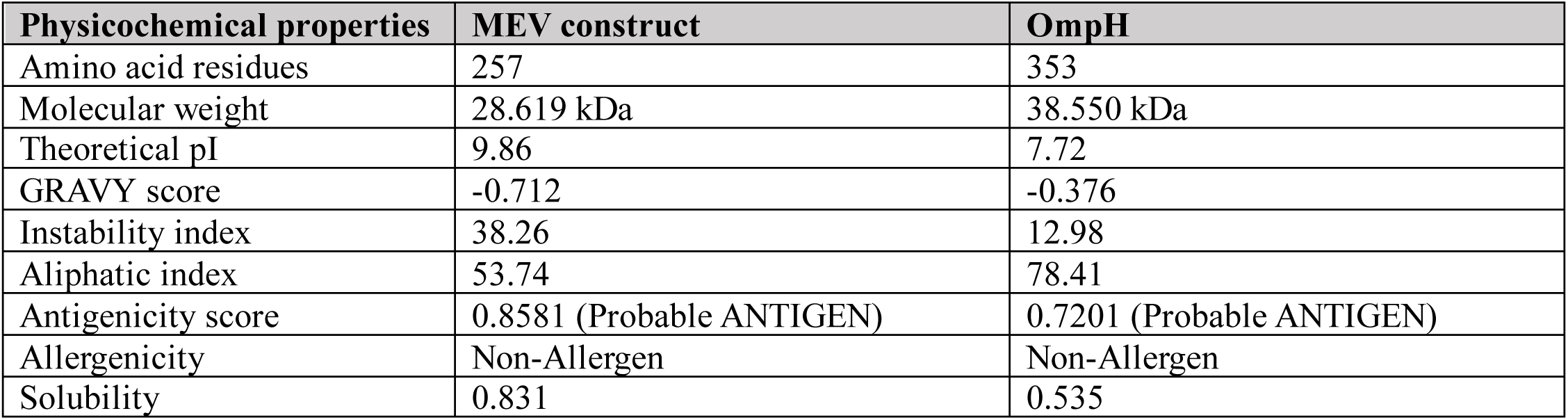
Comparison of physicochemical profile of the designed MEV with OmpH (positive control).

### 3.6 Prediction of secondary and tertiary structures of the MEV construct

The secondary structure analysis using three prediction tools indicated a high proportion of random coils in MEV (Figures 5b and S3), which suggested an enhanced potential of antigenic epitope presentation ^108^. Because the designed MEV includes epitopes derived from intrinsically flexible ECL regions and lacks a suitable experimental structural template, the predicted tertiary model showed relatively low confidence (pTM = 0.22; pLDDT = 45.64). The model was further optimized using the GalaxyRefine server, and the best-refined model was selected based on the lowest clash and MolProbity scores, and the highest GDT-HA (Global Distance Test - High Accuracy) score. This model underwent a second round of refinement and was selected for further analysis (Figure S4). The refined MEV structure was visualized using PyMOL, and selected surface-exposed LBE, CTL, and HTL epitopes were mapped onto the tertiary structure (Figure 5c).

### 3.7 Model quality assessment of MEV construct

The quality of the model was assessed using PROCHECK, which generated a Ramachandran plot (Figure 5d). Analysis revealed that 97.2% of the amino acid residues fell within the favored regions, 2.8% in additionally allowed regions, and no residue in disallowed region. To assess the accuracy of the predicted structure, the quality of the model was further evaluated using ProSA-webserver, which showed a Z-score of −0.98 (Figure 5e), consistent with experimentally determined structures. Overall, the predicted tertiary structure of the MEV indicated that it is a good vaccine design.

### 3.8 Molecular docking with human TLR4 and TLR2

Molecular docking was performed to predict potential interactions between the MEV construct (ligand molecule) and human immune receptors - TLR4 and TLR2, based on energy minimization and structural complementarity at the receptor binding site. Multiple docking clusters were generated, and the top-ranked cluster was selected for further analysis. The best-docked complex of MEV-TLR4 and MEV-TLR2 showed docking scores of −414.16 and −362.40, respectively, with confidence scores of 0.9 (Figures 6 and S5). PRODIGY server predicted favorable binding interactions for MEV-TLR4 (ΔG = −23.3 kcal/mol; K_d_= 7.7 × 10^−18^ M) and MEV-TLR2 (−20.7 kcal/mol; K_d_= 6.3 × 10^−16^ M) complexes, comparable to those predicted for the corresponding OmpH-TLR4 (ΔG = −19.9 kcal/mol; K_d_= 2.4 × 10^−15^ M), and OmpH-TLR2 (ΔG = −18.0 kcal/mol; K_d_= 6.1 × 10^−14^ M) complexes (Table S6). Furthermore, the PDBePISA analysis revealed that the MEV formed 12 hydrogen bonds with TLR4 and its coreceptor MD-2 (Figure 6a) and 13 hydrogen bonds with TLR2 within the range of 5 Å interatomic distance (Figure 6b).

**Fig. 6.**
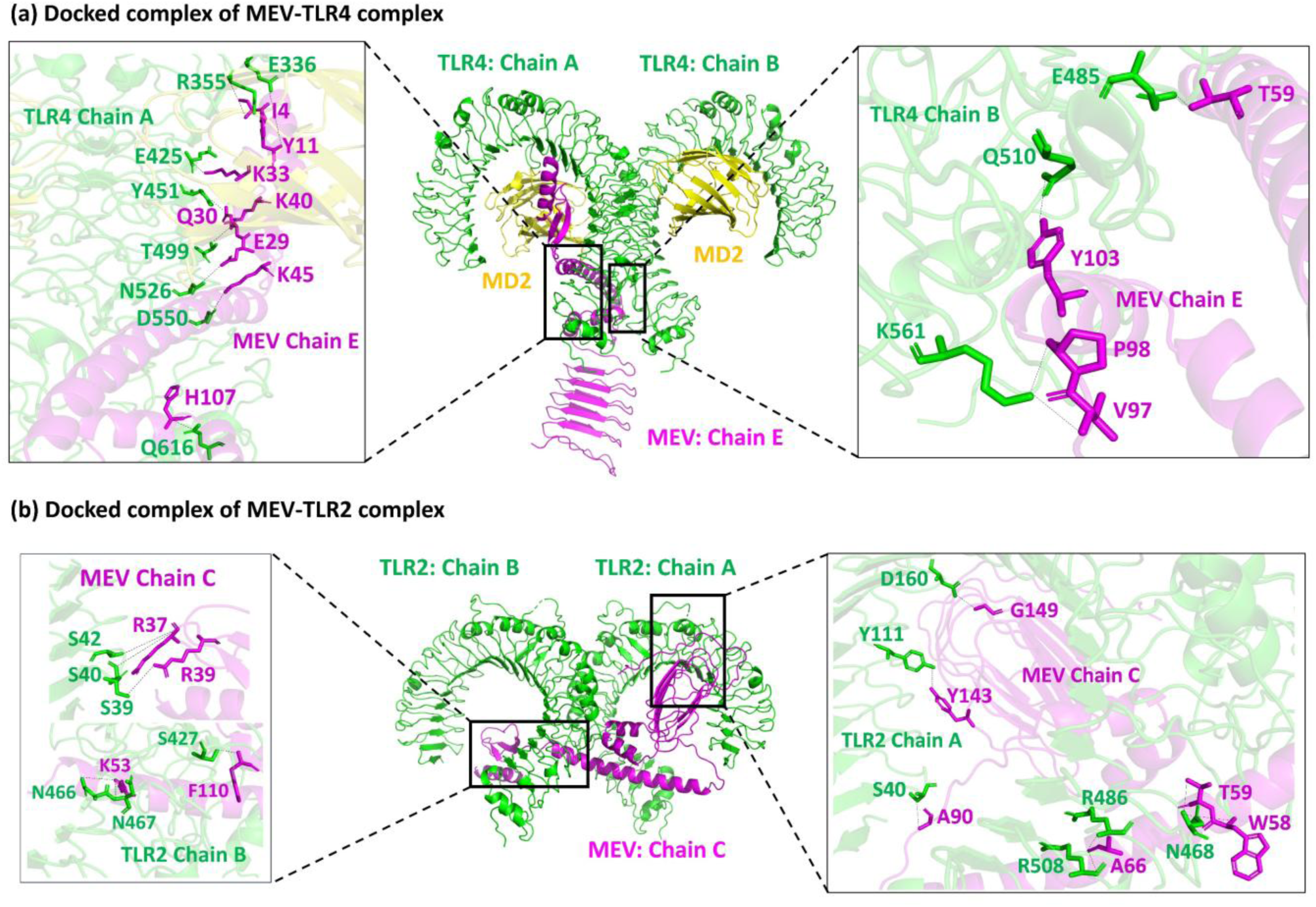
Molecular docking of the MEV construct with TLR4 and TLR2. **(a)** The docked Complex of the vaccine construct with TLR4 (PDB Id: 3FXI), along with the co-receptor myeloid differentiation factor 2 (MD2). The binding interactions between the vaccine construct and TLR4 were analyzed using the PDBePISA server. **(b)** Docked complex of the MEV construct with TLR2 (PDB Id: 6NIG), along with binding interactions analyzed using PDBePISA server.

### 3.9 Residue flexibility analysis and NMA of MEV-TLR4/TLR2 complexes

CABS-flex predicted residue-wise RMSF values for MEV-TLR4/TLR2 complexes over a time period equivalent to 10 ns simulation. Higher RMSF values indicate greater flexibility, while lower values reflect restricted motion of the system. RMSF values for MEV-TLR4 complex (0.04-5.65 Å) (Figure 7a) and MEV-TLR2 complex (0.1-7.23 Å) (Figure 7b) indicated relatively low residue-level fluctuations, with localized increases in flexibility at the MEV-TLR4/TLR2 docking interface attributable to induced-fit conformational adjustments (Figure 7). Moreover, medoids RMSD values (representative structures of each cluster) for MEV-TLR4 complex (2.3-3.3 Å) and MEV-TLR2 complex (4.0-4.8 Å) relative to the input reference structure suggested limited structural variation throughout the simulation, supporting the overall structural stability of both complexes. The positive control also showed a comparable RMSF profile for OmpH-TLR2 complex (0.04-7.18 Å), whereas OmpH-TLR4 complex exhibited higher fluctuations (0.07-10.82 Å) (Figure S6), suggesting our MEV showed comparable residue-level conformational behavior with OmpH.

**Fig. 7.**
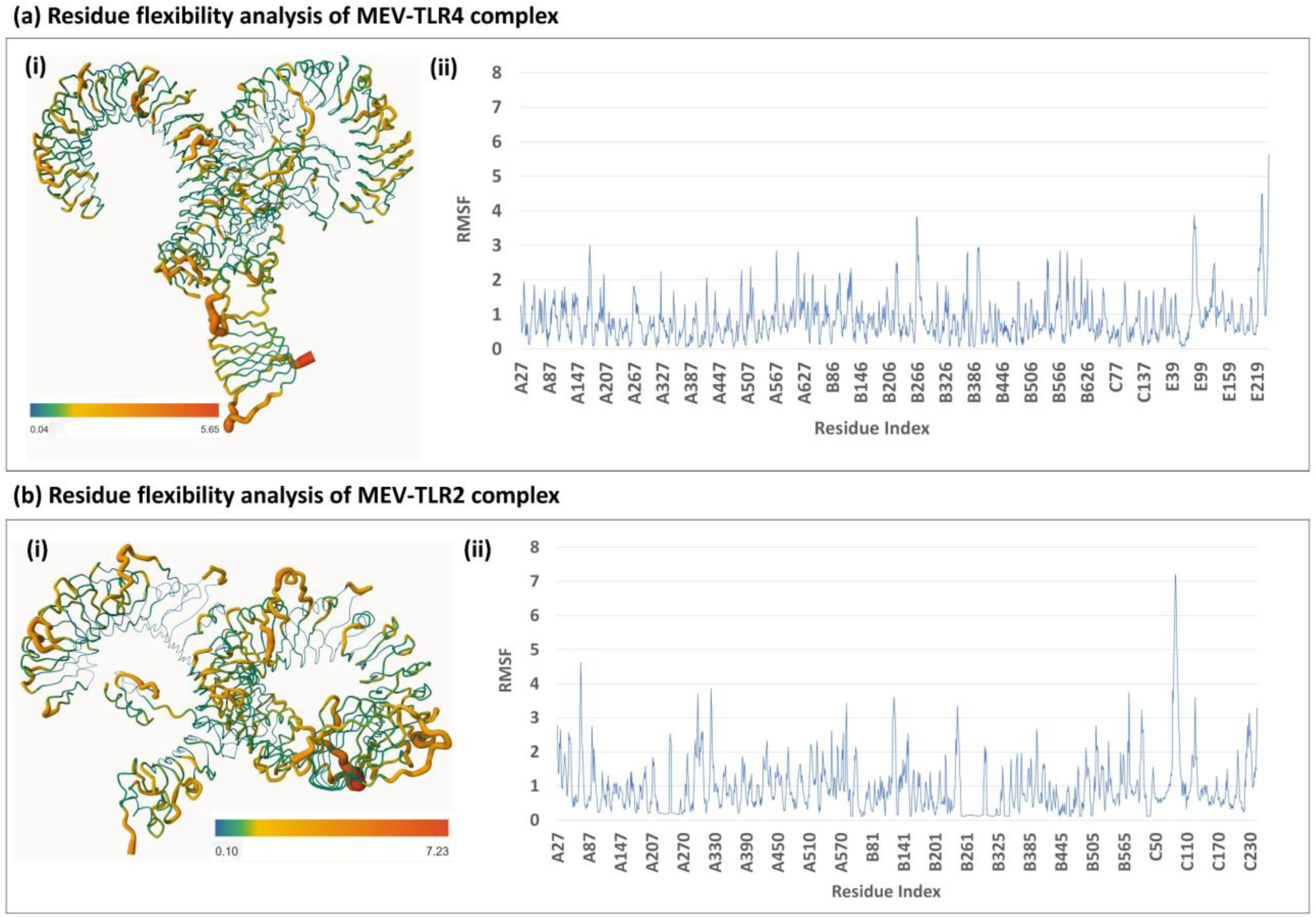
Residue flexibility analysis of MEV-TLR4 and MEV-TLR2 complexes. **(a)** MEV-TLR4 complex and **(b)** MEV-TLR2 complex. **(i)** Structural representation of residue flexibility in the protein-ligand complex. **(ii)** RMSF profiles of the corresponding complexes throughout the simulation. Blue and red regions indicate stable residues and highly flexible residues, respectively.

NMA was further performed using iMODS to characterize the low-frequency modes and flexible conformational states accessible to the MEV-TLR4 and MEV-TLR2 complexes around their equilibrium positions ^109^. The analysis generated elastic network, covariance, B-factor, eigenvalue, deformability, and variance profiles (Figures 8 and S7). The MEV-TLR4/2 covariance maps showed predominantly positively correlated motions along the diagonal (red regions), which indicated coordinated movements among neighboring regions, with localized anti-correlated motions (blue regions). The variance profiles for both complexes demonstrated that conformational fluctuations were distributed across multiple modes rather than being dominated by a single mode, while the elastic network models revealed extensive interatomic contacts (Figure 8a, b). Both complexes exhibited low deformability across most regions, with localized flexibility confined to termini and loop regions, consistent with the B-factor profiles (Figure S6). The estimated eigenvalues for MEV-TLR4 (2.97 × 10^−^⁶) and MEV-TLR2 (1.68 × 10^−^⁵) were comparable to those of the corresponding OmpH complexes (TLR4, 1.59 × 10^−^⁵; TLR2, 2.56 × 10^−^⁵) that suggested greater adaptability of the MEV-receptor complexes. Overall, the NMA profiles of the MEV-immune receptor complexes were comparable to those of the positive control (Figure S8).

**Fig. 8.**
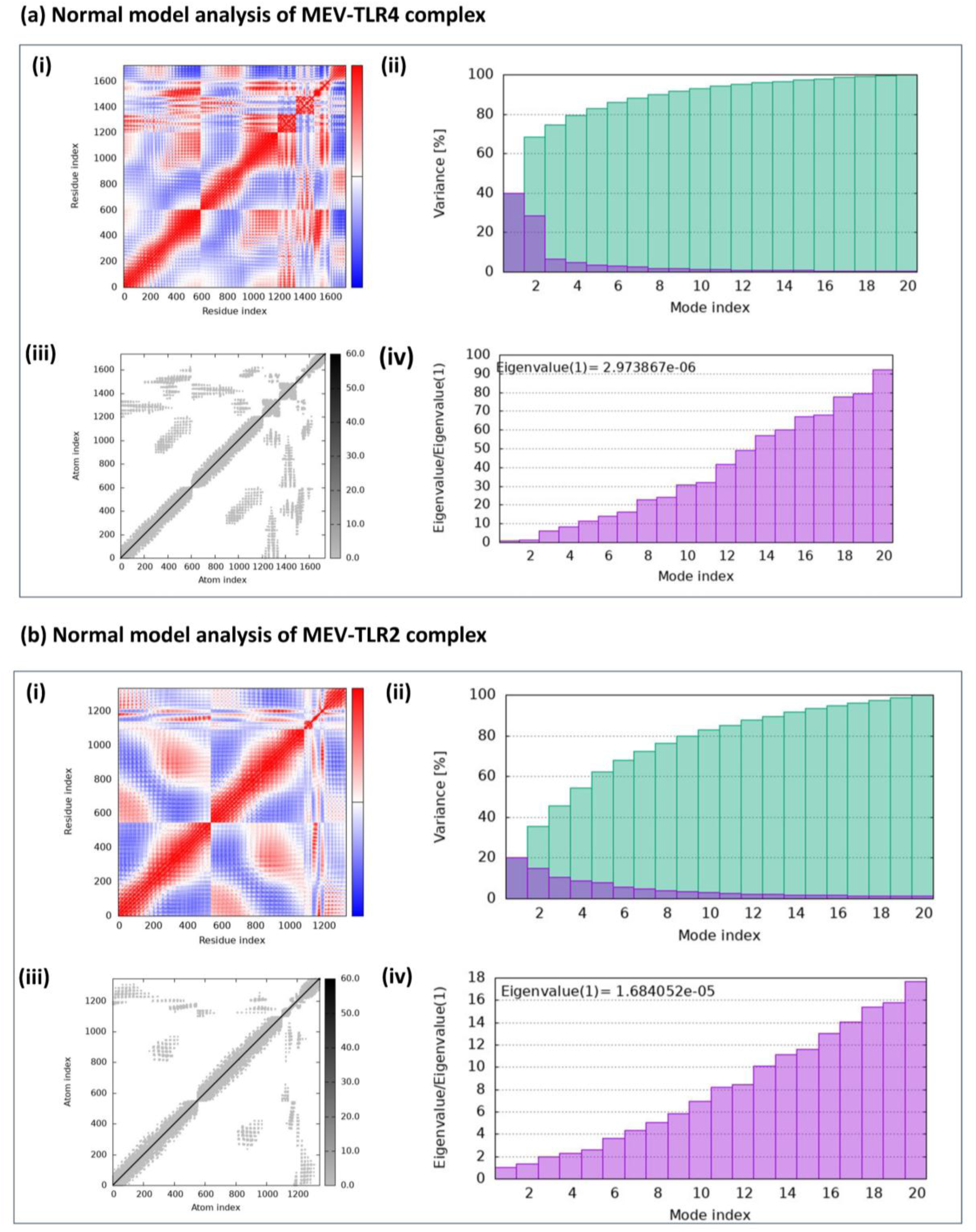
Normal mode analysis of MEV-TLR4 and MEV-TLR2 complexes. **(a)** MEV-TLR4 complex and **(b)** MEV-TLR2 complex. **(i)** Covariance map **(ii)** Variance plot **(iii)** Elastic network model **(iv)** Eigenvalue spectra

### 3.10 MD simulation of MEV-TLR4/TLR2 complexes

The RMSD profiles of the MEV-TLR4/TLR2 and OmpH-TLR4/TLR2 complexes were analyzed over a period of 100 ns MD simulation using Desmond module in the Schrodinger suite.

#### MEV-TLR4 complex

The TLR4 backbone RMSD initially increased (∼ 2.2 Å-4.5 Å) during the first 30 ns, followed by a decrease to 2.7 Å for the remainder of the simulation. In contrast, the MEV RMSD increased continuously (∼3.4-20 Å) during the first 50 ns and subsequently stabilized at 20-21 Å. The initial fluctuations in the MEV RMSD indicated conformational flexibility of the construct, followed by stabilization without evident dissociation (Figure 9a). The OmpH-TLR4 complex showed an initial increase in TLR4 RMSD followed by stabilization at ∼4-5 Å, while OmpH fluctuated around ∼7-8 Å during the later stages of the simulation (Figure 9b). The higher RMSD observed for MEV relative to OmpH indicated greater conformational rearrangements; however, the subsequent plateau phase suggested attainment of a relatively stable conformational state towards the end of the simulation.

**Fig. 9.**
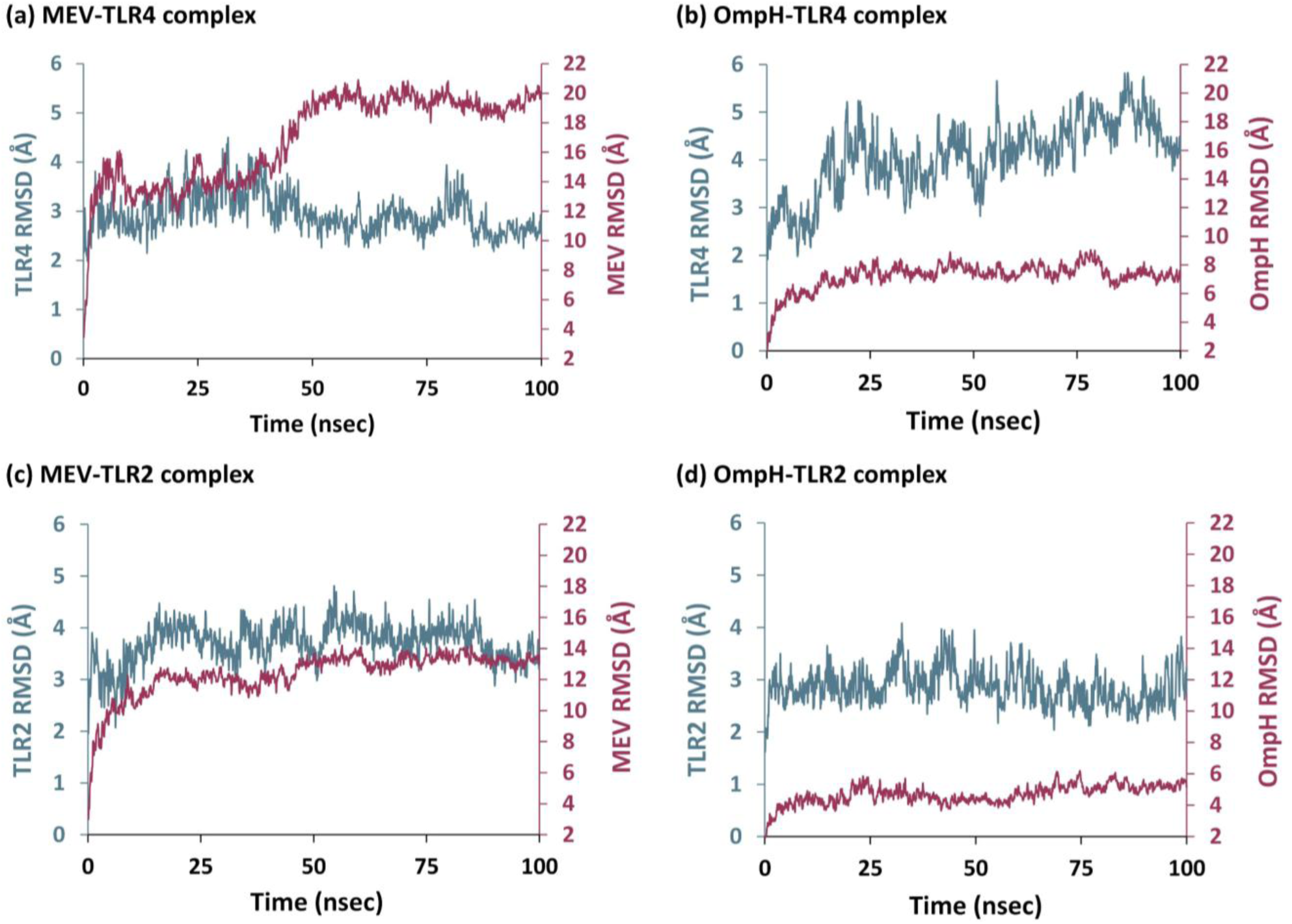
RMSD plots of MEV-TLR4/2 and OmpH-TLR4/2 complexes over 100 ns MD simulation. RMSD profiles for **(a)** MEV-TLR4, **(b)** OmpH-TLR4, **(c)** MEV-TLR2, and **(d)** OmpH-TLR2 complexes.

#### MEV-TLR2 complex

The TLR2 RMSD increased rapidly from ∼2.5 Å to ∼3.8 Å within the first few ns and remained relatively stable (average RMSD ∼3.70 Å) with minimal fluctuations throughout the simulation. The MEV RMSD increased (∼3.8 Å to ∼12.5 Å) during the initial phase (first 15 ns) and then remained comparatively stable with an average RMSD of 12.34 Å (Figure 9c). In OmpH-TLR2 complex, the TLR2 backbone exhibited a similar pattern of initial conformational adjustment followed by stabilization (Figure 9d), while OmpH showed an average RMSD of 3.13 Å throughout the simulation.

Overall, MD simulations of MEV-TLR4 and MEV-TLR2 complexes exhibited relatively stable plateau phases, comparable to OmpH-TLR4/2 complexes, suggesting conformational stability without evident ligand dissociation during the simulation.

### 3.11 Immune simulation of the MEV construct

Immune simulation was performed to evaluate the predicted immunological response elicited by the MEV construct (Figure 10). Following the primary dose, IgM levels initially increased but declined later due to class switching to IgG1 and IgG2 (Figure 10a). Upon subsequent exposures, the levels of IgM + IgG, IgG1 + IgG2, IgG1, and IgG2 were markedly higher than those observed in the primary response, which reflected a strong secondary immune response. The B-cell population increased after each vaccine administration, which suggested long-term immune memory (Figure 10b). Vaccination induced an increase in HTL population (Figure 10c), where TH1 cells dominated the early immune response and accounting for nearly 100% of the TH cell population (Figure 10e). The simulated CTL population remained relatively stable (near the baseline) over the simulated period (Figure 10d). Cytokines IFN-γ and IL-2 were produced at substantially higher concentrations compared to other cytokines (Figure 10f), suggesting that the induced immunity is TH1-driven. The positive control OmpH displayed a similar immune profile, with few notable differences, including a slightly higher antibody and lower TH-cell responses (Figure S9). Overall, the MEV elicited an immune-response pattern comparable to that of the established OmpH antigen.

**Fig. 10.**
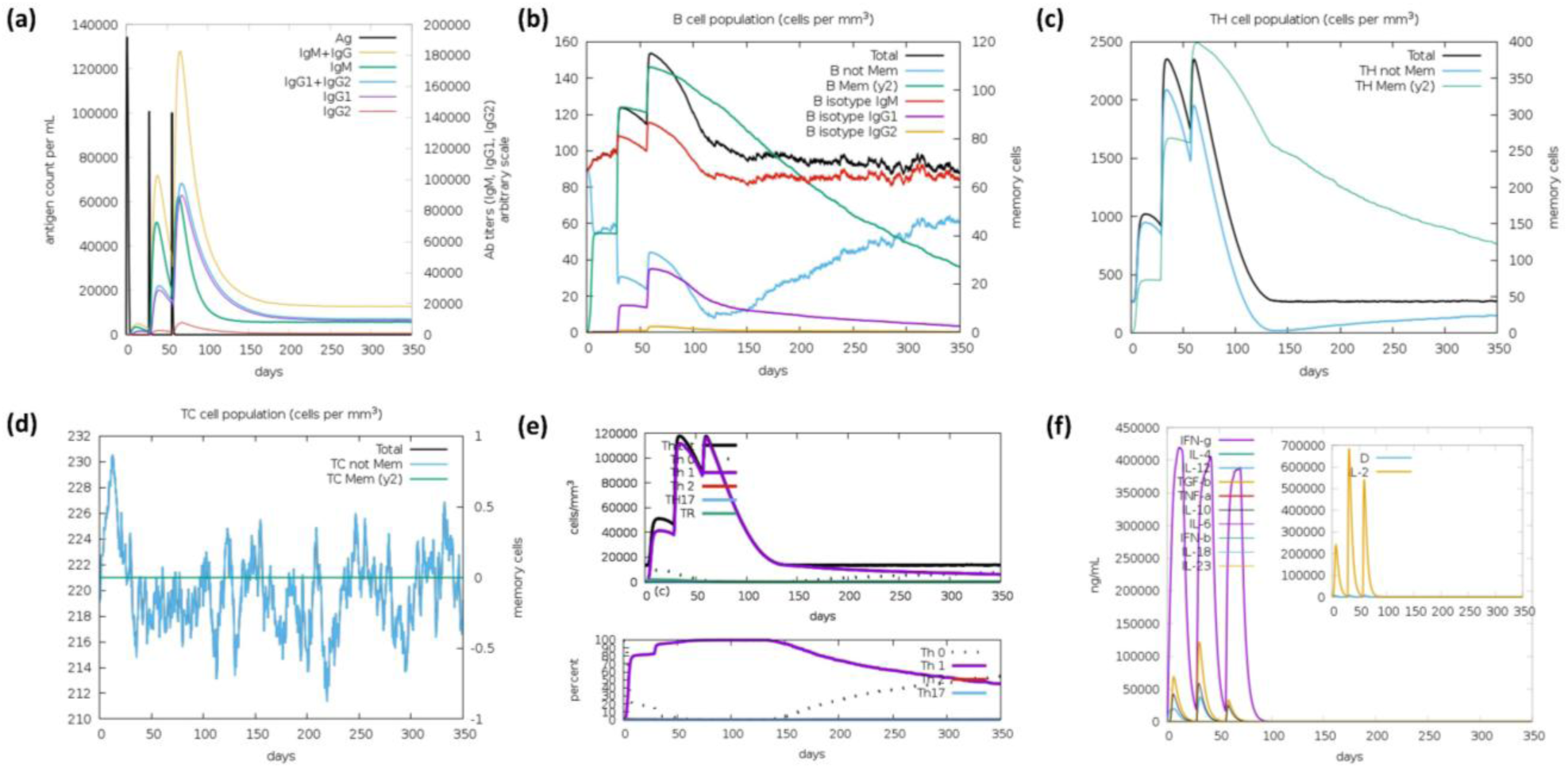
In silico immune simulation of the MEV construct. **(a)** Antigen and subtypes of immunoglobulin levels. **(b)** B-cell isotypes in various states. **(c)** Total helper T-cell population **(d)** Cytotoxic T-cell population **(e)** Helper T-cell isotypes in various states. **(f)** Concentration of cytokines and interleukins.

### 3.12 Codon optimization and in silico cloning

The Pm MEV gene sequence was optimized to enhance its expression in the *E. coli* K-12 expression system. The MEV gene sequence was 771 bp in length, with an estimated CAI of 0.94. This indicated a high level of compatibility with the codon usage bias of the host. The GC content was 50.97%, which falls within the optimal range (40-60%) for efficient transcription and translation. The optimized gene was subsequently in silico cloned into the pET28a (+) expression vector using Benchling. The recombinant construct contained the MEV gene inserted downstream of the T7 promoter with a C-terminal His-tag (Figure 11).

**Fig. 11.**
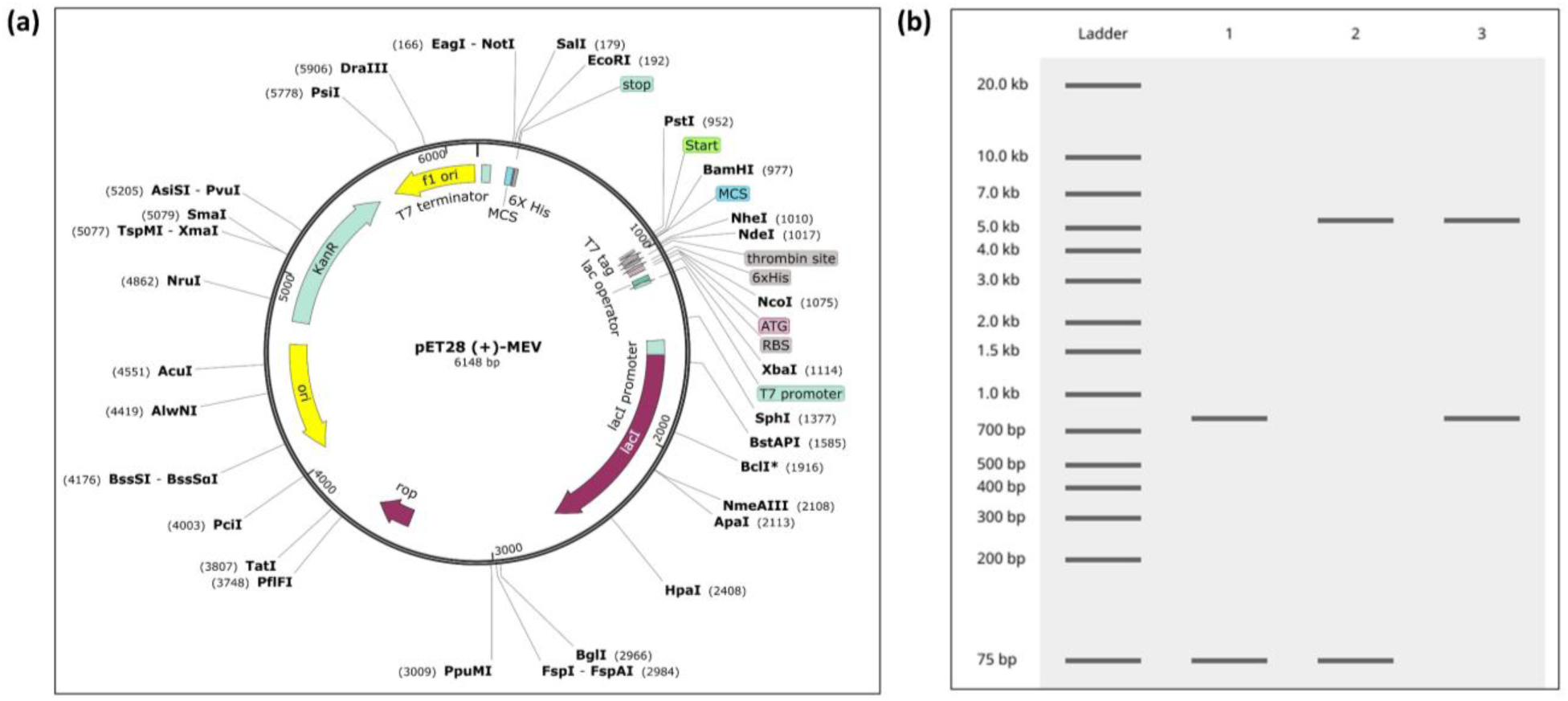
In-silico cloning of the codon-optimized MEV construct into the pET-28a (+) expression vector. **(a)** Schematic representation of the recombinant pET 28a plasmid showing the optimized MEV gene inserted into the multiple cloning site (MCS) downstream of the T7 promoter. **(b)** In silico restriction digestion of the recombinant plasmid showing the expected DNA fragments. Lane 1: MEV construct PCR products; Lane 2: pET-28a vector, and Lane 3: Recombinant plasmid.

## 4. Discussion

Pasteurellosis caused by Pm affects a wide range of animals, including companion animals (cats and dogs), livestock and poultry, resulting in substantial economic losses ^1^. Pm infection can manifest as skin and soft tissue infections, endocarditis, septicemia, meningitis, urinary tract infections, and respiratory infections. In this study, we systematically identified and characterized OMBB proteins in Pm and utilized them as potential vaccine candidates. Using a consensus-based computational framework, we identified 29 OMBB proteins and classified them into diverse functional groups, including TonB-dependent receptors, porins, OM assembly proteins, autotransporters, enzymatic OMPs, secretion system components, adhesins, and efflux proteins. Notably, our OMBB screening pipeline identified four well-known β-barrel protective antigens (OmpH: WP_324023090.1; OmpA: WP_046333314.1; Omp87: WP_083004621.1; and OmpW: WP_083005347.1). The identification of these known OMBB proteins within our dataset underscores the robustness of our designed pipeline. In contrast, TonB-dependent receptor-domain containing proteins and efflux pumps have been described in Pm; however, their structures and precise functions in Past9 remain uncharacterized. The remaining proteins are primarily annotated in the NCBI database with no experimental evidence supporting their characterization. Structural validation using five independent modeling tools supported the prediction of β-barrel architecture, and the resulting models, when structurally aligned, indicated minimal structural deviation (RMSD values < 6 Å). Amino acid sequence variations were distributed throughout the proteins, including ECL, TM, and ICL regions.

Since OMBB proteins are in direct contact with the host, they can serve as promising candidates for MEV design. Although MEV constructs and vaccine candidates have previously been proposed against pasteurellosis for livestock and poultry ^9,26,27^, no licensed human vaccine is currently available. Surface-exposed epitopes from OMBB proteins are highly accessible to the host immune system ^16^, and several OMP-based subunit vaccines have demonstrated protective efficacy in animal models of pasteurellosis ^22–25^. However, antigen accessibility and immunogenicity do not necessarily imply protection as recombinant Pm OmpA has been reported to elicit a strong but non-protective Th2 response in mice model ^110^. Therefore, we applied multiple screening criteria to the identified proteins and selected antigenic, non-allergenic, non-toxic, conserved, and surface-exposed B-cell and T-cell epitopes that were non-homologous to the human proteome. The identification of conserved epitopes is intended to support cross-strain protection, although no cross-strain challenge was performed. Non-homologous epitopes reduce the risk of unintended cross-reactivity with the human proteins. The proposed MEV construct includes epitopes from seven OMBB proteins identified through systematic screening of the Pm proteome (2241 proteins), rather than from a limited number of pre-selected antigens. This distinguishes our approach from earlier proposed MEV designs ^26,27,111^. The selected epitopes were assembled into a 257 amino acid long MEV construct, and its immunogenicity was enhanced by adding two adjuvants-β-defensin 3 and PADRE peptides. The construct exhibited favorable physicochemical properties, high antigenicity, and good solubility in *E. coli* for efficient recombinant production. Molecular docking between the MEV model and TLR4/TLR2 predicted favorable interactions, primarily mediated by adjuvants, facilitating initial receptor engagement, while the incorporated epitopes may contribute to subsequent antigen-specific recognition. Furthermore, MD simulation (100 ns) supported overall conformational stability of both MEV-TLR4/TLR2 complexes. The docking and simulation results were comparable with those obtained for the positive control (OmpH), suggesting that the proposed construct exhibits a similar interaction profile and structural stability. Furthermore, MEV elicited an immune-response pattern comparable to that of the established OmpH antigen. The successful in silico cloning of the MEV construct into pET 28a (+) expression vector supported the feasibility of the construct for future recombinant protein expression and experimental evaluation. Even though our in silico predictions are promising, experimental validation remains essential to confirm them, and therefore, we propose the construct as a candidate for further testing rather than as a validated immunogen.

## 5. Conclusion

This study proposed an MEV construct utilizing surface-exposed Pm OMBB proteins against human pasteurellosis. The designed MEV construct exhibited favorable physicochemical and immunological properties along with structural stability. The vaccine construct revealed stable interactions with human TLR4/2. Although our in silico analyses present a promising vaccine candidate, future studies will require in vitro and in vivo validation, followed by pharmacological evaluation, to confirm its immunogenicity, protective efficacy, and safety.

## Acknowledgements

AP is a recipient of senior research fellowship from the Department of Biotechnology (Grant number: DBT/2023-24/UOD/2326), Government of India. JK is a recipient of junior research fellowship from the University Grants Commission (Reference number: 231620067077), Government of India.

## Author contributions statement

**Amisha Panda:** methodology, software, investigation, data analysis, data curation, writing—original draft preparation. **Jahnvi Kapoor:** data analysis, visualization, writing—reviewing and editing. **Sanjiv Kumar:** conceptualization, methodology, software, writing—reviewing and editing, supervision. **Anannya Bandyopadhyay**: methodology, data analysis, writing—original draft preparation, writing—reviewing and editing, supervision. All authors reviewed the final draft of the manuscript. All authors contributed to the manuscript revision, read and approved the submitted version.

## Data Availability Statement

The data supporting the findings of this study have been deposited in the Zenodo public repository, accessible at https://doi.org/10.5281/zenodo.22741115.

## References

1. Harper M, Boyce JD, Adler B. Pasteurella multocida pathogenesis: 125 years after Pasteur. FEMS Microbiol Lett. 2006;265(1):1–10. doi:10.1111/j.1574-6968.2006.00442.x

2. Giordano A, Dincman T, Clyburn BE, Steed LL, Rockey DC. Clinical Features and Outcomes of Pasteurella multocida Infection. Medicine (Baltimore). 2015;94(36):e1285. doi:10.1097/MD.0000000000001285

3. Hasani SJ, Enferadi A, Sarani S, Nofouzi K. A review of pasteurellosis in humans and animals. Journal of Zoonotic Diseases. 2025;9(3):838–851. doi:10.22034/jzd.2024.18077

4. Iaria C, Cascio A. Please, do not forget Pasteurella multocida. Clin Infect Dis. 2007;45(7):940. doi:10.1086/521247

5. Cuevas I, Carbonero A, Cano D, García-Bocanegra I, Amaro MÁ, Borge C. Antimicrobial resistance of Pasteurella multocida type B isolates associated with acute septicemia in pigs and cattle in Spain. BMC Vet Res. 2020;16(1):222. doi:10.1186/s12917-020-02442-z

6. Kristinsson G. Pasteurella multocida infections. Pediatr Rev. 2007;28(12):472–473. doi:10.1542/pir.28-12-472

7. Oh YH, Moon DC, Lee YJ, Hyun BH, Lim SK. Antimicrobial resistance of Pasteurella multocida strains isolated from pigs between 2010 and 2016. Vet Rec Open. 2018;5(1):e000293. doi:10.1136/vetreco-2018-000293

8. Bae HG, Sohn JH, Seo H, et al. Antimicrobial Susceptibility of Human Pasteurella multocida Isolates in Korea, 2022-2025. Infect Chemother. 2026;58(2):237–242. doi:10.3947/ic.2026.0009

9. Li Y, Xiao J, Chang YF, et al. Immunogenicity and protective efficacy of the recombinant Pasteurella multocida lipoproteins VacJ and PlpE, and outer membrane protein H from P. multocida A:1 in ducks. Front Immunol. 2022;13:985993. doi:10.3389/fimmu.2022.985993

10. St Michael F, Li J, Vinogradov E, Larocque S, Harper M, Cox AD. Structural analysis of the lipopolysaccharide of Pasteurella multocida strain VP161: identification of both Kdo-P and Kdo-Kdo species in the lipopolysaccharide. Carbohydr Res. 2005;340(1):59–68. doi:10.1016/j.carres.2004.10.017

11. Costerton JW, Ingram JM, Cheng KJ. Structure and function of the cell envelope of gram-negative bacteria. Bacteriol Rev. 1974;38(1):87–110. doi:10.1128/br.38.1.87-110.1974

12. Rimler RB, Rhoades KR. Serogroup F, a new capsule serogroup of Pasteurella multocida. J Clin Microbiol. 1987;25(4):615–618. doi:10.1128/jcm.25.4.615-618.1987

13. Carter GR. Studies on Pasteurella multocida. I. A hemagglutination test for the identification of serological types. Am J Vet Res. 1955;16(60):481–484. https://www.ncbi.nlm.nih.gov/pubmed/13238757

14. Carter GR. Further observations on typing Pasteurella multocida by the indirect hemagglutination test. Can J Comp Med Vet Sci. 1962;26(10):238–240. https://www.ncbi.nlm.nih.gov/pubmed/17649399

15. Heddleston KL, Gallagher JE, Rebers PA. Fowl cholera: Gel diffusion precipitin test for serotyping Pasteurella multocida from avian species. Avian Dis. 1972;16(4):925. doi:10.2307/1588773

16. Koebnik R, Locher KP, Van Gelder P. Structure and function of bacterial outer membrane proteins: barrels in a nutshell. Mol Microbiol. 2000;37(2):239–253. doi:10.1046/j.1365-2958.2000.01983.x

17. Fairman JW, Noinaj N, Buchanan SK. The structural biology of β-barrel membrane proteins: a summary of recent reports. Curr Opin Struct Biol. 2011;21(4):523–531. doi:10.1016/j.sbi.2011.05.005

18. Horne JE, Brockwell DJ, Radford SE. Role of the lipid bilayer in outer membrane protein folding in Gram-negative bacteria. J Biol Chem. 2020;295(30):10340–10367. doi:10.1074/jbc.REV120.011473

19. Solan R, Pereira J, Lupas AN, Kolodny R, Ben-Tal N. Gram-negative outer-membrane proteins with multiple β-barrel domains. Proc Natl Acad Sci U S A. 2021;118(31):e2104059118. doi:10.1073/pnas.2104059118

20. E-komon T, Burchmore R, Herzyk P, Davies R. Predicting the outer membrane proteome of Pasteurella multocida based on consensus prediction enhanced by results integration and manual confirmation. BMC Bioinformatics. 2012;13(1):63. doi:10.1186/1471-2105-13-63

21. Wheeler R. Outer membrane proteomics of Pasteurella multocida isolates to identify putative host-specificity determinants. Biosci Horiz Int J Stud Res. 2009;2(1):1–12. doi:10.1093/biohorizons/hzp002

22. Okay S, Özcengiz E, Gürsel I, Özcengiz G. Immunogenicity and protective efficacy of the recombinant Pasteurella lipoprotein E and outer membrane protein H from Pasteurella multocida A:3 in mice. Res Vet Sci. 2012;93(3):1261–1265. doi:10.1016/j.rvsc.2012.05.011

23. Varinrak T, Poolperm P, Sawada T, Sthitmatee N. Cross-protection conferred by immunization with an rOmpH-based intranasal fowl cholera vaccine. Avian Pathol. 2017;46(5):515–525. doi:10.1080/03079457.2017.1321105

24. Kumar A, Yogisharadhya R, Ramakrishnan MA, Viswas KN, Shivachandra SB. Structural analysis and cross-protective efficacy of recombinant 87 kDa outer membrane protein (Omp87) of Pasteurella multocida serogroup B:2. Microb Pathog. 2013;65:48–56. doi:10.1016/j.micpath.2013.09.007

25. Shivachandra SB, Kumar A, Yogisharadhya R, Viswas KN. Immunogenicity of highly conserved recombinant VacJ outer membrane lipoprotein of Pasteurella multocida. Vaccine. 2014;32(2):290–296. doi:10.1016/j.vaccine.2013.10.075

26. Zhang R, Dai L, Jia Y, et al. Evaluation of a multi-epitope vaccine PME for Pasteurella multocida in mouse model. Front Immunol. 2025;16(1652907):1652907. doi:10.3389/fimmu.2025.1652907

27. Qiao Y, Wu A, Li H, et al. Design of a multi-Epitope vaccine against Ovine Pasteurella multocida using immunoinformatics strategies. Microorganisms. 2026;14(3):656. doi:10.3390/microorganisms14030656

28. Kapoor J, Panda A, Naqvi I, Ganta S, Kumar S, Bandyopadhyay A. Identification and characterization of outer membrane proteins and membrane spanning protein complexes in Brucella melitensis. Proteins. 2026;(prot.70118). doi:10.1002/prot.70118

29. Panda A, Kapoor J, Hareramadas B, et al. Identification and characterization of novel outer membrane proteins of Brachyspira pilosicoli. Microbe Immun. 2025;2(4):79. doi:10.36922/mi025230050

30. Panda A, Kapoor J, Hareramadas B, et al. Design of a Multi-Epitope Vaccine using β-barrel Outer Membrane Proteins Identified in Chlamydia trachomatis. J Membr Biol. Published online September 4, 2025. doi:10.1007/s00232-025-00360-5

31. Smallman TR, Perlaza-Jiménez L, Wang X, et al. Pathogenomic analysis and characterization of Pasteurella multocida strains recovered from human infections. Microbiol Spectr. 2024;12(4):e0380523. doi:10.1128/spectrum.03805-23

32. Sayers EW, Bolton EE, Brister JR, et al. Database resources of the National Center for Biotechnology Information in 2023. Nucleic Acids Res. 2023;51(D1):D29–D38. doi:10.1093/nar/gkac1032

33. Rice P, Longden I, Bleasby A. EMBOSS: The European molecular biology open software suite. Trends Genet. 2000;16(6):276–277. doi:10.1016/s0168-9525(00)02024-2

34. Sachdeva G, Kumar K, Jain P, Ramachandran S. SPAAN: a software program for prediction of adhesins and adhesin-like proteins using neural networks. Bioinformatics. 2005;21(4):483–491. doi:10.1093/bioinformatics/bti028

35. Almagro Armenteros JJ, Tsirigos KD, Sønderby CK, et al. SignalP 5.0 improves signal peptide predictions using deep neural networks. Nat Biotechnol. 2019;37(4):420–423. doi:10.1038/s41587-019-0036-z

36. Yu CS, Lin CJ, Hwang JK. Predicting subcellular localization of proteins for Gram-negative bacteria by support vector machines based on n-peptide compositions. Protein Sci. 2004;13(5):1402–1406. doi:10.1110/ps.03479604

37. Yu NY, Wagner JR, Laird MR, et al. PSORTb 3.0: improved protein subcellular localization prediction with refined localization subcategories and predictive capabilities for all prokaryotes. Bioinformatics. 2010;26(13):1608–1615. doi:10.1093/bioinformatics/btq249

38. Roumia AF, Tsirigos KD, Theodoropoulou MC, Tamposis IA, Hamodrakas SJ, Bagos PG. OMPdb: A global hub of beta-barrel outer membrane proteins. Front Bioinform. 2021;1:646581. doi:10.3389/fbinf.2021.646581

39. Hallgren J, Tsirigos KD, Pedersen MD, et al. DeepTMHMM predicts alpha and beta transmembrane proteins using deep neural networks. *bioRxiv*. Published online April 10, 2022. doi:10.1101/2022.04.08.487609

40. Ou YY, Gromiha MM, Chen SA, Suwa M. TMBETADISC-RBF: Discrimination of beta-barrel membrane proteins using RBF networks and PSSM profiles. Comput Biol Chem. 2008;32(3):227–231. doi:10.1016/j.compbiolchem.2008.03.002

41. Bernhofer M, Rost B. TMbed: transmembrane proteins predicted through language model embeddings. BMC Bioinformatics. 2022;23(1):326. doi:10.1186/s12859-022-04873-x

42. Abramson J, Adler J, Dunger J, et al. Accurate structure prediction of biomolecular interactions with AlphaFold 3. Nature. 2024;630(8016):493–500. doi:10.1038/s41586-024-07487-w

43. Lin Z, Akin H, Rao R, et al. Evolutionary-scale prediction of atomic-level protein structure with a language model. Science. 2023;379(6637):1123–1130. doi:10.1126/science.ade2574

44. Waterhouse A, Bertoni M, Bienert S, et al. SWISS-MODEL: homology modelling of protein structures and complexes. Nucleic Acids Res. 2018;46(W1):W296–W303. doi:10.1093/nar/gky427

45. Baek M, DiMaio F, Anishchenko I, et al. Accurate prediction of protein structures and interactions using a three-track neural network. Science. 2021;373(6557):871–876. doi:10.1126/science.abj8754

46. Du Z, Su H, Wang W, et al. The trRosetta server for fast and accurate protein structure prediction. Nat Protoc. 2021;16(12):5634–5651. doi:10.1038/s41596-021-00628-9

47. Schrödinger, L., & DeLano, W. (2020). PyMOL.

48. Zhang C, Shine M, Pyle AM, Zhang Y. US-align: universal structure alignments of proteins, nucleic acids, and macromolecular complexes. Nat Methods. 2022;19(9):1109–1115. doi:10.1038/s41592-022-01585-1

49. Kapoor J, Panda A, Rajagopal R, Kumar S, Bandyopadhyay A. Structural characterization of the type IV secretion system in *Brucella melitensis* for virtual screening-based therapeutic targeting. J Chem Inf Model. 2026;(acs.jcim.6c00870). doi:10.1021/acs.jcim.6c00870

50. Sievers F, Wilm A, Dineen D, et al. Fast, scalable generation of high-quality protein multiple sequence alignments using Clustal Omega. Mol Syst Biol. 2011;7(1):539. doi:10.1038/msb.2011.75

51. Ponomarenko J, Bui HH, Li W, et al. ElliPro: a new structure-based tool for the prediction of antibody epitopes. BMC Bioinformatics. 2008;9(1):514. doi:10.1186/1471-2105-9-514

52. Vita R, Blazeska N, Marrama D, et al. The Immune Epitope Database (IEDB): 2024 update. Nucleic Acids Res. 2025;53(D1):D436–D443. doi:10.1093/nar/gkae1092

53. Reynisson B, Alvarez B, Paul S, Peters B, Nielsen M. NetMHCpan-4.1 and NetMHCIIpan-4.0: improved predictions of MHC antigen presentation by concurrent motif deconvolution and integration of MS MHC eluted ligand data. Nucleic Acids Res. 2020;48(W1):W449–W454. doi:10.1093/nar/gkaa379

54. Altschul SF, Gish W, Miller W, Myers EW, Lipman DJ. Basic local alignment search tool. J Mol Biol. 1990;215(3):403–410. doi:10.1016/S0022-2836(05)80360-2

55. Marrama D, Chronister WD, Westernberg L, et al. PEPMatch: a tool to identify short peptide sequence matches in large sets of proteins. BMC Bioinformatics. 2023;24(1):485. doi:10.1186/s12859-023-05606-4

56. Doytchinova IA, Flower DR. VaxiJen: a server for prediction of protective antigens, tumour antigens and subunit vaccines. BMC Bioinformatics. 2007;8(1):4. doi:10.1186/1471-2105-8-4

57. Sharma N, Patiyal S, Dhall A, Pande A, Arora C, Raghava GPS. AlgPred 2.0: an improved method for predicting allergenic proteins and mapping of IgE epitopes. Brief Bioinform. 2021;22(4). doi:10.1093/bib/bbaa294

58. Lomize AL, Todd SC, Pogozheva ID. Spatial arrangement of proteins in planar and curved membranes by PPM 3.0. Protein Sci. 2022;31(1):209–220. doi:10.1002/pro.4219

59. Bui HH, Sidney J, Dinh K, Southwood S, Newman MJ, Sette A. Predicting population coverage of T-cell epitope-based diagnostics and vaccines. BMC Bioinformatics. 2006;7(1):153. doi:10.1186/1471-2105-7-153

60. Dong R, Chu Z, Yu F, Zha Y. Contriving multi-Epitope subunit of vaccine for COVID-19: Immunoinformatics approaches. Front Immunol. 2020;11:1784. doi:10.3389/fimmu.2020.01784

61. Mortazavi B, Molaei A, Fard NA. Multi-epitopevaccines, from design to expression; an in silico approach. Hum Immunol. 2024;85(3):110804. doi:10.1016/j.humimm.2024.110804

62. Chen Z, Zhu Y, Sha T, et al. Design of a new multi-epitope vaccine against Brucella based on T and B cell epitopes using bioinformatics methods. Epidemiol Infect. 2021;149(e136):e136. doi:10.1017/S0950268821001229

63. Wilkins MR, Gasteiger E, Bairoch A, et al. Protein identification and analysis tools in the ExPASy server. Methods Mol Biol. 1999;112:531–552. https://www.ncbi.nlm.nih.gov/pubmed/10027275

64. Hon J, Marusiak M, Martinek T, et al. SoluProt: prediction of soluble protein expression in Escherichia coli. Bioinformatics. 2021;37(1):23–28. doi:10.1093/bioinformatics/btaa1102

65. Jones DT, Cozzetto D. DISOPRED3: precise disordered region predictions with annotated protein-binding activity. Bioinformatics. 2015;31(6):857–863. doi:10.1093/bioinformatics/btu744

66. Tesfaye AB, Werid GM, Tao Z, et al. Advances in Pasteurella multocida vaccine development: From conventional to next-generation strategies. Vaccines (Basel). 2025;13(10):1034. doi:10.3390/vaccines13101034

67. Geourjon C, Deléage G. SOPMA: significant improvements in protein secondary structure prediction by consensus prediction from multiple alignments. Comput Appl Biosci. 1995;11(6):681–684. doi:10.1093/bioinformatics/11.6.681

68. Jones DT. Protein secondary structure prediction based on position-specific scoring matrices. J Mol Biol. 1999;292(2):195–202. doi:10.1006/jmbi.1999.3091

69. Høie MH, Kiehl EN, Petersen B, et al. NetSurfP-3.0: accurate and fast prediction of protein structural features by protein language models and deep learning. Nucleic Acids Res. 2022;50(W1):W510–W515. doi:10.1093/nar/gkac439

70. Heo L, Park H, Seok C. GalaxyRefine: Protein structure refinement driven by side-chain repacking. Nucleic Acids Res. 2013;41(Web Server issue):W384–8. doi:10.1093/nar/gkt458

71. Laskowski RA, MacArthur MW, Thornton JM. PROCHECK: validation of protein-structure coordinates. In: International Tables for Crystallography. International Union of Crystallography; 2012:684–687. doi:10.1107/97809553602060000882

72. Wiederstein M, Sippl MJ. ProSA-web: interactive web service for the recognition of errors in three-dimensional structures of proteins. Nucleic Acids Res. 2007;35(Web Server issue):W407–10. doi:10.1093/nar/gkm290

73. Krissinel E, Henrick K. Inference of macromolecular assemblies from crystalline state. J Mol Biol. 2007;372(3):774–797. doi:10.1016/j.jmb.2007.05.022

74. Vangone A, Bonvin AMJJ. PRODIGY: A contact-based predictor of binding affinity in protein-protein complexes. Bio Protoc. 2017;7(3):e2124. doi:10.21769/BioProtoc.2124

75. Wróblewski K, Zalewski M, Kuriata A, Kmiecik S. CABS-flex 3.0: an online tool for simulating protein structural flexibility and peptide modeling. Nucleic Acids Res. 2025;53(W1):W95–W101. doi:10.1093/nar/gkaf412

76. López-Blanco JR, Aliaga JI, Quintana-Ortí ES, Chacón P. iMODS: internal coordinates normal mode analysis server. Nucleic Acids Res. 2014;42(Web Server issue):W271–6. doi:10.1093/nar/gku339

77. Schrödinger Release 2025-3: Schrödinger Suite, Schrödinger, LLC, New York, NY, 2025.. Schrodinger LLC. 2025. https://www.schrodinger.com/citations/

78. Rapin N, Lund O, Bernaschi M, Castiglione F. Computational immunology meets bioinformatics: the use of prediction tools for molecular binding in the simulation of the immune system. PLoS One. 2010;5(4):e9862. doi:10.1371/journal.pone.0009862

79. Castiglione F, Mantile F, De Berardinis P, Prisco A. How the interval between prime and boost injection affects the immune response in a computational model of the immune system. Comput Math Methods Med. 2012;2012:842329. doi:10.1155/2012/842329

80. Grote A, Hiller K, Scheer M, et al. JCat: a novel tool to adapt codon usage of a target gene to its potential expression host. Nucleic Acids Res. 2005;33(Web Server issue):W526–31. doi:10.1093/nar/gki376

81. Bosch M, Garrido ME, Llagostera M, Pérez De Rozas AM, Badiola I, Barbé J. Characterization of the Pasteurella multocida hgbA gene encoding a hemoglobin-binding protein. Infect Immun. 2002;70(11):5955–5964. doi:10.1128/IAI.70.11.5955-5964.2002

82. Ogunnariwo JA, Schryvers AB. Characterization of a novel transferrin receptor in bovine strains of Pasteurella multocida. J Bacteriol. 2001;183(3):890–896. doi:10.1128/JB.183.3.890-896.2001

83. Shen X, Guan L, Zhang J, Xue Y, Si L, Zhao Z. Study in the iron uptake mechanism of Pasteurella multocida. Vet Res. 2025;56(1):41. doi:10.1186/s13567-025-01469-0

84. Marandi M, Mittal KR. Characterization of an outer membrane protein of Pasteurella multocida belonging to the OmpA family. Vet Microbiol. 1996;53(3-4):303–314. doi:10.1016/s0378-1135(96)01219-9

85. Katoch S, Sharma M, Patil RD, Kumar S, Verma S. In vitro and in vivo pathogenicity studies of Pasteurella multocida strains harbouring different ompA. Vet Res Commun. 2014;38(3):183–191. doi:10.1007/s11259-014-9601-6

86. Dabo SM, Confer AW, Quijano-Blas RA. Molecular and immunological characterization of Pasteurella multocida serotype A:3 OmpA: evidence of its role in P. multocida interaction with extracellular matrix molecules. Microb Pathog. 2003;35(4):147–157. doi:10.1016/s0882-4010(03)00098-6

87. Simms AN, Jerse AE. In vivo selection for Neisseria gonorrhoeae opacity protein expression in the absence of human carcinoembryonic antigen cell adhesion molecules. Infect Immun. 2006;74(5):2965–2974. doi:10.1128/IAI.74.5.2965-2974.2006

88. Cole JG, Fulcher NB, Jerse AE. Opacity proteins increase Neisseria gonorrhoeae fitness in the female genital tract due to a factor under ovarian control. Infect Immun. 2010;78(4):1629–1641. doi:10.1128/IAI.00996-09

89. Yogisharadhya R, Mohanty NN, Chacko N, Mondal M, Chanda MM, Shivachandra SB. Sequence and structural analysis of OmpW protein of Pasteurella multocida strains reveal evolutionary conservation among members of Pasteurellaceae along with its homologues. Gene Rep. 2019;14:36–44. doi:10.1016/j.genrep.2018.11.004

90. Grover M. Conservation of properties of outer membranes protein across host genera of Pasteurella multocida suggests common mechanism of action. Mol Biol (Los Angel). 2016;5(2). doi:10.4172/2168-9547.1000161

91. Konovalova A, Kahne DE, Silhavy TJ. Outer membrane biogenesis. Annu Rev Microbiol. 2017;71:539–556. doi:10.1146/annurev-micro-090816-093754

92. Qiao S, Luo Q, Zhao Y, Zhang XC, Huang Y. Structural basis for lipopolysaccharide insertion in the bacterial outer membrane. Nature. 2014;511(7507):108–111. doi:10.1038/nature13484

93. Botte M, Ni D, Schenck S, et al. Cryo-EM structures of a LptDE transporter in complex with Pro-macrobodies offer insight into lipopolysaccharide translocation. Nat Commun. 2022;13(1):1826. doi:10.1038/s41467-022-29459-2

94. Goh KJ, Stubenrauch CJ, Lithgow T. The TAM, a Translocation and Assembly Module for protein assembly and potential conduit for phospholipid transfer. EMBO Rep. 2024;25(4):1711–1720. doi:10.1038/s44319-024-00111-y

95. Harris JA, Roy K, Woo-Rasberry V, et al. Directed evaluation of enterotoxigenic Escherichia coli autotransporter proteins as putative vaccine candidates. PLoS Negl Trop Dis. 2011;5(12):e1428. doi:10.1371/journal.pntd.0001428

96. Mizan S, Henk A, Stallings A, Maier M, Lee MD. Cloning and characterization of sialidases with 2-6’ and 2-3’ sialyl lactose specificity from Pasteurella multocida. J Bacteriol. 2000;182(24):6874–6883. doi:10.1128/JB.182.24.6874-6883.2000

97. Braun V, Ondraczek R, Hobbie S. Activation and secretion of Serratia hemolysin. Zentralbl Bakteriol. 1993;278(2-3):306–315. doi:10.1016/s0934-8840(11)80847-9

98. Guérin J, Bigot S, Schneider R, Buchanan SK, Jacob-Dubuisson F. Two-partner secretion: Combining efficiency and simplicity in the secretion of large proteins for bacteria-host and bacteria-bacteria interactions. Front Cell Infect Microbiol. 2017;7:148. doi:10.3389/fcimb.2017.00148

99. Delattre AS, Clantin B, Saint N, Locht C, Villeret V, Jacob-Dubuisson F. Functional importance of a conserved sequence motif in FhaC, a prototypic member of the TpsB/Omp85 superfamily. FEBS J. 2010;277(22):4755–4765. doi:10.1111/j.1742-4658.2010.07881.x

100. Kaur D, Gandhi S, Mukhopadhaya A. Salmonella Typhimurium adhesin OmpV activates host immunity to confer protection against systemic and gastrointestinal infection in mice. Infect Immun. 2021;89(8):e0012121. doi:10.1128/IAI.00121-21

101. Hatfaludi T, Al-Hasani K, Dunstone M, Boyce J, Adler B. Characterization of TolC efflux pump proteins from Pasteurella multocida. Antimicrob Agents Chemother. 2008;52(11):4166–4171. doi:10.1128/AAC.00245-08

102. MacRaild CA, Seow J, Das SC, Norton RS. Disordered epitopes as peptide vaccines. Pept Sci (Hoboken). 2018;110(3):e24067. doi:10.1002/pep2.24067

103. Ferris LK, Mburu YK, Mathers AR, et al. Human beta-defensin 3 induces maturation of human langerhans cell-like dendritic cells: an antimicrobial peptide that functions as an endogenous adjuvant. J Invest Dermatol. 2013;133(2):460–468. doi:10.1038/jid.2012.319

104. Ghaffari-Nazari H, Tavakkol-Afshari J, Jaafari MR, Tahaghoghi-Hajghorbani S, Masoumi E, Jalali SA. Improving multi-Epitope long peptide vaccine potency by using a strategy that enhances CD4+ T help in BALB/c mice. PLoS One. 2015;10(11):e0142563. doi:10.1371/journal.pone.0142563

105. Baseer S, Ahmad S, Ranaghan KE, Azam SS. Towards a peptide-based vaccine against Shigella sonnei: A subtractive reverse vaccinology based approach. Biologicals. 2017;50:87–99. doi:10.1016/j.biologicals.2017.08.004

106. Vessely C, Estey T, Randolph TW, et al. Stability of a trivalent recombinant protein vaccine formulation against botulinum neurotoxin during storage in aqueous solution. J Pharm Sci. 2009;98(9):2970–2993. doi:10.1002/jps.21498

107. Shone C, Agostini H, Clancy J, et al. Bivalent recombinant vaccine for botulinum neurotoxin types A and B based on a polypeptide comprising their effector and translocation domains that is protective against the predominant A and B subtypes. Infect Immun. 2009;77(7):2795–2801. doi:10.1128/IAI.01252-08

108. Yu M, Zhu Y, Li Y, et al. Design of a novel multi-Epitope vaccine against Echinococcus granulosus in immunoinformatics. Front Immunol. 2021;12:668492. doi:10.3389/fimmu.2021.668492

109. A. Bauer J, Bauerová-Hlinková V. Normal mode analysis: A tool for better understanding protein flexibility and dynamics with application to homology models. In: Homology Molecular Modeling - Perspectives and Applications. IntechOpen; 2021. doi:10.5772/intechopen.94139

110. Dabo SM, Confer A, Montelongo M, York P, Wyckoff JH 3rd. Vaccination with Pasteurella multocida recombinant OmpA induces strong but non-protective and deleterious Th2-type immune response in mice. Vaccine. 2008;26(34):4345–4351. doi:10.1016/j.vaccine.2008.06.029

111. Geng B, Zhou B, Zhang Z, Jiang G, Zhu W. No more than three PlpE non-overlapping epitopes trigger significant antibody production in individuals vaccinated with the Pasteurella multocida epitope-chimeric proteins. Microbiol Spectr. 2026;14(3):e0287825. doi:10.1128/spectrum.02878-25

