## Supplemental material for "Characterization of Outer Membrane β-barrel Proteins of *Pasteurella multocida* for Multi-Epitope Vaccine Design against Human Pasteurellosis"

**Supplemental Tables**

**Table S1:** Comprehensive results from computational tools employed in our study to predict outer membrane β-barrel proteins.

| **Protein accession no.** | **Protein annotation (NCBI)^≠^** | **Pepstats** | | **SPAAN** | **SignalP** | | **PSORTb** | | **CELLO** | **OMPdb^$^** | **TMBETADISC-RBF** | **DeepTMHMM** | **TMbed^&^** | |
| --- | --- | --- | --- | --- | --- | --- | --- | --- | --- | --- | --- | --- | --- | --- |
| **GROUP A** |  | **AA** | **MolWt (kDa)** |  | **Prediction** | **Cleavage site prediction (residue number)** | **Localization** | **Score^*^** |  |  |  |  | **TMbed_b** | **TMbed_B** |
| WP_096743251.1 | TonB-dependent receptor domain-containing protein | 726 | 81.2 | 0.361947 | SP(Sec/SPI) | 22-23 | Outer Membrane | 10 | Outer  Membrane | 500 | Outer Membrane Protein | BETA | 97 | 90 |
| WP_104228050.1 | TonB-dependent receptor plug domain-containing protein | 803 | 92.4 | 0.127665 | SP(Sec/SPI) | 26-27 | Outer Membrane | 9.52 | Outer  Membrane | 223 | Outer Membrane Protein | BETA | 99 | 91 |
| WP_126417535.1 | TonB-dependent hemoglobin/transferrin/lactoferrin family receptor | 784 | 89.7 | 0.409363 | SP(Sec/SPI) | 26-27 | Outer Membrane | 10 | Outer  Membrane | 500 | Outer Membrane Protein | BETA | 99 | 94 |
| WP_249984729.1 | TonB-dependent hemoglobin/transferrin/lactoferrin family receptor | 989 | 113.8 | 0.291661 | SP(Sec/SPI) | 21-22 | Outer Membrane | 10 | Outer  Membrane | 724 | Outer Membrane Protein | BETA | 94 | 92 |
| WP_249984736.1 | TonB-dependent hemoglobin/transferrin/lactoferrin family receptor | 967 | 110.3 | 0.203452 | SP(Sec/SPI) | 24-25 | Outer Membrane | 10 | Outer  Membrane | 778 | Outer Membrane Protein | BETA | 94 | 88 |
| WP_046333314.1 | Porin OmpA | 354 | 38 | 0.614961 | SP(Sec/SPI) | 21-22 | Outer Membrane | 10 | Outer  Membrane | 500 | Outer Membrane Protein | BETA | 41 | 36 |
| WP_249984777.1 | Opacity family porin | 198 | 21.8 | 0.248443 | SP(Sec/SPI) | 20-21 | Outer Membrane | 10 | Outer  Membrane | 229 | Outer Membrane Protein | BETA | 37 | 34 |
| WP_324023089.1 | Porin | 345 | 38.3 | 0.516594 | SP(Sec/SPI) | 20-21 | Outer Membrane | 10 | Outer  Membrane | 500 | Outer Membrane Protein | BETA | 70 | 65 |
| WP_324023090.1 | Porin | 342 | 37 | - | SP(Sec/SPI) | 20-21 | Outer Membrane | 10 | Outer  Membrane | 500 | Outer Membrane Protein | BETA | 67 | 65 |
| WP_083004621.1 | Outer membrane protein assembly factor BamA | 791 | 87.7 | 0.226478 | SP(Sec/SPI) | 18-19 | Outer Membrane | 10 | Outer  Membrane | 500 | Outer Membrane Protein | BETA | 74 | 71 |
| WP_126417754.1 | LPS assembly protein LptD | 787 | 91.2 | 0.351811 | SP(Sec/SPI) | 29-30 | Outer Membrane | 10 | Outer  Membrane | 500 | Outer Membrane Protein | BETA | 116 | 97 |
| WP_249984417.1 | Surface lipoprotein assembly modifier | 506 | 58.3 | 0.256227 | SP(Sec/SPI) | 29-30 | Unknown | 2 | Outer  Membrane | 500 | Outer Membrane Protein | BETA | 66 | 64 |
| WP_324022736.1 | Autotransporter assembly complex family protein | 586 | 67.1 | 0.167425 | SP(Sec/SPI) | 30-31 | Outer Membrane | 10 | Outer  Membrane | 373 | Outer Membrane Protein | BETA | 74 | 70 |
| WP_249984510.1 | Autotransporter domain-containing protein | 679 | 74.5 | 0.344184 | SP(Sec/SPI) | 20-21 | Unknown | 7 | Outer  Membrane | 4 | Outer Membrane Protein | BETA | 53 | 52 |
| WP_249984830.1 | Autotransporter outer membrane beta-barrel domain-containing protein | 850 | 95 | 0.397918 | SP(Sec/SPI) | 24-25 | Unknown | 7 | Outer  Membrane | 162 | Outer Membrane Protein | BETA | 52 | 51 |
| WP_115299510.1 | Sialidase family protein | 1060 | 118 | 0.308431 | SP(Sec/SPI) | 25-26 | Outer Membrane | 9.83 | Outer  Membrane | 2 | Outer Membrane Protein | BETA | 55 | 51 |
| WP_324022762.1 | Phospholipase A | 290 | 33.8 | 0.301686 | SP(Sec/SPI) | 29-30 | Outer Membrane | 10 | Outer  Membrane | 500 | Outer Membrane Protein | BETA | 55 | 47 |
| WP_083004763.1 | ShlB/FhaC/HecB family hemolysin secretion/activation protein | 576 | 64.6 | 0.280238 | SP(Sec/SPI) | 30-31 | Outer Membrane | 10 | Outer  Membrane | 500 | Outer Membrane Protein | BETA | 77 | 69 |
| WP_324023075.1 | ShlB/FhaC/HecB family hemolysin secretion/activation protein | 618 | 69.7 | 0.329116 | SP(Sec/SPI) | 18-19 | Outer Membrane | 9.99 | Outer  Membrane | 500 | Outer Membrane Protein | BETA | 75 | 71 |
| WP_249984768.1 | MipA/OmpV family protein | 257 | 28.8 | 0.17828 | SP(Sec/SPI) | 21-22 | Outer Membrane | 10 | Outer  Membrane | 69 | Outer Membrane Protein | BETA | 55 | 44 |
| WP_108511463.1 | Outer membrane protein transport protein | 436 | 46.9 | 0.332066 | SP(Sec/SPI) | 25-26 | Outer Membrane | 10 | Outer  Membrane | 500 | Outer Membrane Protein | BETA | 64 | 57 |
| WP_005757734.1 | Outer membrane beta-barrel protein | 232 | 26.5 | 0.112022 | SP(Sec/SPI) | 18-19 | Unknown | 2.5 | Outer Membrane | 27 | Outer Membrane Protein | BETA | 37 | 36 |
| **GROUP B** |  |  |  |  |  |  |  |  |  |  |  |  |  |  |
| WP_249984435.1 | TonB-dependent receptor | 805 | 91.2 | 0.18053 | SP(Sec/SPI) | 26-27 | Outer Membrane | 9.49 | Outer  Membrane | 503 | Non-Outer Membrane Protein | BETA | 99 | 91 |
| WP_249984789.1 | TonB-dependent receptor | 797 | 89.5 | 0.388649 | SP(Sec/SPI) | 22-23 | Outer Membrane | 9.52 | Outer Membrane | 504 | Non-Outer Membrane Protein | BETA | 99 | 92 |
| WP_324023022.1 | TonB-dependent receptor domain-containing protein | 883 | 99.8 | 0.070765 | SP(Sec/SPI) | 27-28 | Cytoplasmic | 8.96 | Outer Membrane | 500 | Non-Outer Membrane Protein | BETA | 101 | 94 |
| WP_005756417.1 | Toxin/drug exporter Tde | 455 | 50.6 | 0.229879 | LIPO(Sec/SPII) | 18-19 | Outer Membrane | 10 | Outer  Membrane | 500 | Outer Membrane Protein | SP | 20 | 20 |
| WP_104227660.1 | Efflux transporter outer membrane subunit | 463 | 51.9 | 0.237084 | LIPO(Sec/SPII) | 16-17 | Outer Membrane | 10 | Outer  Membrane | 500 | Outer Membrane Protein | SP | 19 | 18 |
| WP_083005347.1 | OmpW family outer membrane protein | 204 | 21.9 | 0.175327 | SP(Sec/SPI) | 21-22 | Outer Membrane | 10 | Periplasmic | 500 | Non-Outer Membrane Protein | BETA | 38 | 36 |
| WP_249984615.1 | Sialidase family protein | 802 | 89.8 | 0.471348 | SP(Sec/SPI) | 21-22 | Periplasmic | 9.44 | Outer  Membrane |  | Outer Membrane Protein | BETA | 54 | 42 |

≠Proteins were named as per their annotation in the NCBI database, searched using protein accession number.

*****A PSORTb score ≥ 7.5 indicates reliable subcellular localization prediction.

$Number of significant matches to outer membrane β-barrel proteins in the OMPdb database.

&**TMbed_B:** Number of transmembrane β-strand oriented from the inside to outside of the membrane; **TMbed_b:** Number of transmembrane β-strand oriented from the outside to inside of the membrane.

**Table S2:** Confidence scores of OMBB protein structural models predicted from five tools, along with alignment RMSD values obtained via US-align.

| **Protein accession no.** | **Protein annotation (NCBI)^≠^** | **Confidence Scores^†^** | | | | | **Structural alignment (RMSD in Å)** |
| --- | --- | --- | --- | --- | --- | --- | --- |
|  |  | **AlphaFold 3** | **RoseTTAFold** | **TrRosetta** | **ESMFold** | **SWISS-MODEL** |  |
| **GROUP A** |  |  |  |  |  |  |  |
| WP_096743251.1 | TonB-dependent receptor domain-containing protein | 0.92 | 0.803 | 0.926 | 0.925 | 0.87 | 2.14 |
| WP_104228050.1 | TonB-dependent receptor plug domain-containing protein | 0.93 | 0.81 | 0.908 | 0.912 | 0.87 | 3.72 |
| WP_126417535.1 | TonB-dependent hemoglobin/transferrin/lactoferrin family receptor | 0.92 | 0.83 | 0.927 | 0.905 | 0.89 | 3.10 |
| WP_249984729.1 | TonB-dependent hemoglobin/transferrin/lactoferrin family receptor | 0.92 | 0.75 | 0.827 | 0.741 | 0.89 | 4.13 |
| WP_249984736.1 | TonB-dependent hemoglobin/transferrin/lactoferrin family receptor | 0.92 | 0.67 | 0.833 | 0.758 | 0.87 | 4.71 |
| WP_046333314.1 | Porin OmpA | 0.62 | 0.78 | 0.782 | 0.708 | 0.80 | 4.13 |
| WP_249984777.1 | Opacity family porin | 0.78 | 0.86 | 0.922 | 0.911 | 0.84 | 2.53 |
| WP_324023089.1 | Porin | 0.91 | 0.74 | 0.883 | 0.912 | 0.91 | 2.05 |
| WP_324023090.1 | Porin | 0.9 | 0.74 | 0.908 | 0.906 | 0.94 | 2.33 |
| WP_083004621.1 | Outer membrane protein assembly factor BamA | 0.82 | 0.85 | 0.850 | 0.768 | 0.89 | 5.18 |
| WP_126417754.1 | LPS assembly protein LptD | 0.91 | 0.80 | 0.942 | 0.863 | 0.89 | 2.44 |
| WP_249984417.1 | Surface lipoprotein assembly modifier | 0.79 | 0.78 | 0.805 | 0.875 | 0.87 | 2.65 |
| WP_324022736.1 | Autotransporter assembly complex family protein | 0.88 | 0.84 | 0.872 | 0.882 | 0.91 | 3.50 |
| WP_249984510.1 | Autotransporter domain-containing protein | 0.93 | 0.80 | 0.942 | 0.917 | 0.93 | 2.34 |
| WP_249984830.1 | Autotransporter outer membrane beta-barrel domain-containing protein | 0.85 | 0.78 | 0.847 | 0.845 | 0.90 | 2.64 |
| WP_115299510.1 | Sialidase family protein | 0.85 | 0.73 | 0.666 | 0.562 | 0.86 | 5.28 |
| WP_324022762.1 | Phospholipase A | 0.9 | 0.76 | 0.934 | 0.873 | 0.88 | 1.63 |
| WP_083004763.1 | ShlB/FhaC/HecB family hemolysin secretion/activation protein | 0.88 | 0.81 | 0.872 | 0.870 | 0.87 | 2.46 |
| WP_324023075.1 | ShlB/FhaC/HecB family hemolysin secretion/activation protein | 0.91 | 0.80 | 0.931 | 0.871 | 0.88 | 2.37 |
| WP_249984768.1 | MipA/OmpV family protein | 0.9 | 0.74 | 0.904 | 0.907 | 0.89 | 1.68 |
| WP_108511463.1 | Outer membrane protein transport protein | 0.91 | 0.81 | 0.954 | 0.929 | 0.91 | 1.56 |
| WP_005757734.1 | Outer membrane beta-barrel protein | 0.73 | 0.69 | 0.835 | 0.883 | 0.82 | 3.22 |
| **GROUP B** |  |  |  |  |  |  |  |
| WP_249984435.1 | TonB-dependent receptor | 0.92 | 0.76 | 0.867 | 0.866 | 0.85 | 3.13 |
| WP_249984789.1 | TonB-dependent receptor | 0.94 | 0.75 | 0.881 | 0.885 | 0.88 | 3.20 |
| WP_324023022.1 | TonB-dependent receptor domain-containing protein | 0.91 | 0.74 | 0.842 | 0.824 | 0.87 | 3.32 |
| WP_005756417.1 | Toxin/drug exporter Tde | 0.92 | 0.84 | 0.897 | 0.907 | 0.93 | 3.13 |
| WP_104227660.1 | Efflux transporter outer membrane subunit | 0.91 | 0.81 | 0.894 | 0.950 | 0.88 | 3.03 |
| WP_083005347.1 | OmpW family outer membrane protein | 0.83 | 0.76 | 0.948 | 0.902 | 0.91 | 1.27 |
| WP_249984615.1 | Sialidase family protein | 0.92 | 0.75 | 0.631 | 0.548 | 0.92 | 4.91 |

≠Proteins were named as per their annotation in the NCBI database, searched using protein accession number.

**†**AlphaFold 3 pTM score > 0.5 indicates a generally reliable predicted structural model. RoseTTAFold confidence score with values above 0.7-0.8 suggests high model accuracy. In TrRosetta, TM-score evaluates global structural similarity between predicted and native structures; scores above 0.5 imply correct topology. ESMFold pLDDT score ranges from 0 to 1 where higher pLDDT scores (closer to 1.0) indicate higher confidence in the accuracy of the predicted structure. GMQE score in SWISS-MODEL estimates overall model quality based on alignment and template structure; values above 0.6-0.7 indicate good reliability.

**Table S3:** Pm strains used for analysis of sequence variation in the identified OMBB proteins.

| **Assembly accession** | **Assembly name** | **Strain name** | **Genome size (Mb)** | **Scaffolds** | **CDS^#^** | **Submission Date** | **Geographic Location** |
| --- | --- | --- | --- | --- | --- | --- | --- |
| GCF_002948995.1 | ASM294899v1 | 161215033201-1 | 2.126 | 1 | 1901 | 19-02-2018 | Netherlands: Hoofddorp |
| GCF_008693845.1 | ASM869384v1 | FDAARGOS_644 | 2.250 | 1 | 2002 | 25-09-2019 | USA: KY |
| GCF_018139065.1 | ASM1813906v1 | HuN001 | 2.287 | 1 | 2064 | 25-04-2021 | China: Hunan |
| GCF_035223645.1 | ASM3522364v1 | Past29 | 2.275 | 2 | 2043 | 05-01-2024 | Australia: Clayton |
| GCF_035224565.1 | ASM3522456v1 | Past9 | 2.419 | 1 | 2242 | 05-01-2024 | Australia: Clayton |
| GCF_035226565.1 | ASM3522656v1 | Past3 | 2.308 | 2 | 2113 | 05-01-2024 | Australia: Clayton |
| GCF_900478175.1 | 33787_E01 | NCTC10382 | 2.314 | 1 | 2064 | 18-06-2018 | - |
| GCF_900638315.1 | 57195_D02 | NCTC11619 | 2.317 | 1 | 2059 | 20-12-2018 | - |

#CDS: Coding Sequences

**Table S4:** Amino acid sequence variations identified in the 29 OMBB proteins.

| **Protein accession no.** | **Protein annotation (NCBI)^≠^** | **Total number** | **Amino acid sequence variations** |
| --- | --- | --- | --- |
| **GROUP A** |  |  |  |
| WP_096743251.1 | TonB-dependent receptor domain-containing protein | 25 | A19, N53, A63, T91, H156, Q160, I182, Q289, I325, N330, A332, T334, H374, Q395, T400, T402, S403, S414, A428, I463, A530, T559, S592, G627, V688 |
| WP_104228050.1 | TonB-dependent receptor plug domain-containing protein | 712 | M1-P4, P7, L8, K10-T12, A14-C17, S19-S23, S25-F37, T39, E40, V42, Y44, Q47-T58, E60-G70, I72, T73, Y75-N79, H81, R83-K98, E100-S103, A107-N123, L125-F132, A135-Y148, F150-M154, S156-E159, Q161-S163, V165, A167-L169, T173, G175-G186, N188-K203, V205-L227, P229, K230, A232-T240, K242-D246, L248-F252, L254, S255, R257, T258, R260-A287, L289-F291, W293-E302, S304-R306, S308-F331, A333-A336, I338-T349, A351, D353-K384, G386, Y387, D389-E402, A404-P407, L410-T414, S416-L419, G421-T465, Y467-L472, I474, E475, L477-S479, K481-P487, V489, F491-N500, I502, F506-L520, L522-R524, Y526, R528-K559, P561, A563-L566, L568-F570, Q572-R575, L577, L578, K580-G582, I584-N588, N590, R591, I593-Y610, S612-N616, S618-T650, R652-D687, I689-R691, G693-T704, S706-I711, A713-D736, G738-R748, Q750, K752-A763, I765, L766, V768, L769, Q771-F803 |
| WP_126417535.1 | TonB-dependent hemoglobin/transferrin/lactoferrin family receptor | 650 | M1-N9, A11-A26, E28, T30, E31, E33, Q34, T36, Q38-R66, T68, T75, D77, G79-N83, L87, K88, F90, M92, E96, D97, G101, S103, I104, V107-S116, Y118-R120, S127, L129-P133, L135-D140, V142, R143, S145, D146, F148-A150, S154, G158, N160-N162, L164-R181, S183, A185-F204, L208-Q212, T214, H216, F218-P235, V238-Y245, A247, M249-Q257, I259-R275, T278, L280-W284, E286-D288, Q290-E292, L294, A296-G330, R332-V379, Q381-I387, Q389-Y408, I410-K420, H422-H432, G434, L435, H439-K443, K445-V469, T471-I489, T491, Y493, V495, A498, S499, M501-S504, T506-S514, P516, S517, K519, P520, K522-Q545, N547, R549, H550, L552-Q577, M579, V580, L582, K584, I587, Y588, V590, L592-N597, D599-G608, K610-K619, K621-S630, L634-G639, D641, E643-E660, R662-G687, Q689-F715, T717-G719, Y721-A731, I733, F737-K739, H742, D745-L747, G749, N751-S754, L756, S758-E762, K764, L766, Q767, Y770, A771, Y776-L780, I782, R783 |
| WP_249984729.1 | TonB-dependent hemoglobin/transferrin/lactoferrin family receptor | 796 | M1, L3-P6, L9, M10, T12-S17, S19-A21, P23-T26, T29, V31, S33-D37, I39-R55, R57-S62, S64, E71, G73, T75-T79, G83-S85, Y87, V89, D93, E94, M100, L104-E117, F119-G121, T128, N130-I134, N136-T141, T143, K144, A146, I149-S151, A155, S159, M161-E163, K165-K181, Y184-N188, Q190-F204, F208-R212, E214, H216, I218-A243, Y246-N253, I255, L257-F259, P261-H265, F267-K362, D364-S367, V369-K371, V373-D400, I402-H407, K409-S412, Q414-C420, I423, C425-F445, D447-S457, K459-G475, Q477-E479, L481-N497, D500-N502, H504-E522, Q524, K526-G528, L530, Y531, K533-V539, K541, G543-R593, S595-I598, V600, K601, N604-A606, Y608-A610, K612-D622, N624, Y625, K629-I633, R635-F638, V640, V642, P643, G645-D670, M672-H692, Y694-I696, L698-R707, Q709-Y712, N714, F716, A718, S721, D722, M726, T727, K729-P737, V739, N740, A743, I745-A752, T754-F767, D770, K772, N773, I775-I787, S789-H792, P794-Y796, N798, Q799, R801, H803, V806, T807, F809, I811-E816, A818-S825, R828-K832, T834-Q836, M840-A848, R852-N857, G859, S861-G868, D870-A877, K880-E883, Y885-Y888, E890, A893-D906, V909-R912, R915-I919, A921-A923, A925-F935, V937, T941, E943, M946, S950, A951, S953, R955-G958, L962, I963, Q965-T967, A969, I971, K972, F981-A985, L987, T988 |
| WP_249984736.1 | TonB-dependent hemoglobin/transferrin/lactoferrin family receptor | 834 | M1-A10, T12-H27, E29-E32, D34, T35, T37, S39-Q68, S70, E77, G79, T81-A85, G89-S91, Y93, I95, D99, E100, A104, T106, V107, L110-E123, F125-G127, T134, N136-I140, T142-K147, A149, K150, A152, V155-V157, S161, A165, L167-E169, K171-A188, S191-F210, I214-K218, H220, H222, L224-A243, Y246-T253, V255, F257-H265, F267-L283, N286, V288-F424, C426-N456, K458-E487, D489-V528, K530-I578, V580-D602, G604, Y605, D609-Q613, E615-H675, Y677-D684, L686-Y695, K697, F699, A701, S704, D705, L707-T710, K712-P720, P722, V723, P726, E728-A735, T737-Q751, K753, R755, H756, I758-Y784, N786, V787, V789-N791, V794, K795, L797, I799-N804, G806-D813, N816-R825, R827-A836, K840-G845, G847, D849-E866, K868-V897, A899-G901, I903-F913, V915, T919-R921, L924, E927-A929, S931, K933-G936, S938, L940-T945, A947, I949, N950, S954, F959-A963, I965, T966 |
| WP_046333314.1 | Porin OmpA | 281 | K3-A7, I10, A11, L13-A105, H107, N109, H110, A112-Y120, V122-S164, V166, A168, G169, E172-V183, Y185-N189, V191-D203, R205-V225, E227-V229, K231, T232, T234, N236, S237, V239, T240, G242, D244, A246, D247, K249-N254, L256, G258-S270, A272, Y276, D283, T296, A298-Y300, V302-K304, Q308, A310-S312, T314, H316, E318, N320-K326, S329, K331, R333, K334, I337, A338, A341-D343, A349-K351, N353, K354 |
| WP_249984777.1 | Opacity family porin | 96 | V5-V7, L10, T12, A15, A16, Y30, S31, L33, E36-S39, S41-A48, K52-F56, S63, L68-H70, E72-T87, S89-P91, K93, Y94, L96, V98-W101, L106, F111-I115, S124, Q125, H127, K129-L132, F134-P137, Y139-Y142, T145, V147, H148, L155, A161, V163, K164, V166-L168, A170, K179-I185 |
| WP_324023089.1 | Porin | 20 | S12, S17, T146, D168, N172, L205, M225, K236-T239, S241, G249-D254, S256, F307 |
| WP_324023090.1 | Porin | 187 | I5, V6, V10-V13, A15, S17-N19, Y24-D27, N34, V37, L40, K42, K43, K45-E47, G49, V52, N54, S59, K61-S63, D65, A74-A76, L78, T82-I104, N106-K111, Y114, A118, Y119, G121, L122, T124, N129, V137, V139, F145-G147, I149, N151, L153, S154, A159, N161, E166, N168, F170, T171, G174, A175, S179, A180, A182, D183, Q185-F193, V195-G197, K202, M203, V206, L210, Q216, Y218-E228, F231, M232, T235, A240-L242, L244, G245, A249, S251, V253-G258, K260, A262, L263, V265, N268, D270-A275, V277, L281, A284, K286-T293, D295, S297-L300, A302, G303, L306, Q309, E311, F313, V314, G316-G319, E321, D323-V327, T329-V334, V336, H341 |
| WP_083004621.1 | Outer membrane protein assembly factor BamA | 20 | T42, D149, T178, E183, G188, A191, D192, S200, V240, A249, T285, S296, V464, S633, E728, A734-V738 |
| WP_126417754.1 | LPS assembly protein LptD | 15 | M1-K6, V47, L86, F247, H397, A412, A463, D619, I697, A742 |
| WP_249984417.1 | Surface lipoprotein assembly modifier | 25 | M2, R15, Q46, A61, F117, V125, E146, A196, Q199, V205, V227, F248, H275, E333, M334, M344, V370, Q396, S416, Q433, T441, R455, V459, V468, N505 |
| WP_324022736.1 | Autotransporter assembly complex family protein | 9 | E112, I119, Q141, R181, N192, R200, A398, V421, S584 |
| WP_249984510.1 | Autotransporter domain-containing protein | 47 | A47, D49, N50, L53, I68, A93, K105, A106, Q109, T110, A112, A138, A140, V164, A210, A225, S227, E247, W254-F256, W259, A260, A263, L283, V309, G328, A348, T352, D402, Q403, W405, N413, G436, A495, D506, T515, A524, Q563, A573, T593, N609, A616, T624, A638, A655, K663 |
| WP_249984830.1 | Autotransporter outer membrane beta-barrel domain-containing protein | 303 | M1-L130, G145, S158-N162, I164, S165, N194, L198, K200, L208, K212, V213, S216-A218, Y221-D226, S230, N232, A246, N250, F251, H256, S259, P262, R264, V266, V277, D287, P289, S290, K313, E315, P322, T352, T367, I372, H373, M400, A406, D408, E423, S479, H499, Q515, H554, I578, V579, V605, A610, F612, T616, N630, H634, L668, S688, V702, Q705, R706, I708, R710, A712, L720, L728, H731, T732, D735, Y738-Q741, N743-N753, A756, K761, V762, T769-F850 |
| WP_115299510.1 | Sialidase family protein | 911 | N5, P6, V8, L11, I13, S15-F35, V37, H40, E42-Q53, S55-V58, I60, W61, S65-R68, K70, G72-A78, K80, Q83-G86, W88-D90, T93-R96, H99, D101, K103, T104, G106, N107, T110-I152, A154-N162, R164-G198, S200, L202-H283, D285, N287, K289, T290, S292-K303, W305-T311, P313, V315, Q318-N321, N323, L325, I328-I343, R345, G347, T350, D352-P357, D359-T382, S384-V387, L389, D390, G392-M398, N400-S402, R404-K411, G413, Y415, I418-K437, S439-N443, E446-F450, G452-E456, S458, G460, D462-D466, K468, F470, G472-E490, Q492-S494, K496-N503, Y505, V506, M510, E512-D515, S517-L520, Y522, N524-T526, Y528, T530-P535, M538-F542, R544-L568, G572-Q579, M581, G582, S584-K607, F609-D636, S638-I641, F643, R644, S646-K675, N677, K679-Y681, H683, A684, E688, I690-L797, Q799-I805, K808, L810-F822, D824-D828, S831-G839, K841-S845, G847-A874, Y876-W879, G881-H891, L893-D896, S898-Q902, K905-D907, E909-M932, A935, S936, T938, M940-L946, A948, D952-Y964, H966-L974, V976, V978, N979, I982-E986, K988-R1001, L1003-D1006, V1008, A1009, Q1012-H1020, S1022-L1029, R1032, F1033, F1035-F1041, H1043-K1047, Q1049, S1050, A1053-H1057, R1059, F1060 |
| WP_324022762.1 | Phospholipase A | 8 | C13, M18, D95, D99, T201, I218, C222, H225 |
| WP_083004763.1 | ShlB/FhaC/HecB family hemolysin secretion/activation protein | 10 | S11, D65, D84, G101, H235, V400, M444, V473, S505, D563 |
| WP_324023075.1 | ShlB/FhaC/HecB family hemolysin secretion/activation protein | 524 | M1, Y3-S5, F7-F11, A13-S15, F17, A18, P20, T21, L23-K29, E32, Q33, D35-Q37, H40-S57, V59-T62, P64-I84, V86, V87, Y89, T91-Q99, S101-R114, S116-A132, Q134-C147, G149-V155, M157, K159-K167, T172-S183, Q185-T189, I191, L192, R194-K208, T210-V225, N227-R229, E232, S234, L235, N237-D245, T247-I250, P252-L259, E261, D263, A265-T277, G279-A283, S285-T288, W291, G293-A305, L308-T311, F313-R331, S333, L336-Y339, S341, V342, W344-A354, N356, D357, V359-S373, S376-T382, S384-F388, A391-R404, S406-N411, E413-L427, Q430-T450, N453-P457, P459-M470, I472, I473, A475-K481, F483-G498, W500-P504, V506-L511, G515, R516, V519, T522, D523, E525-N546, G548-L555, K557, S561-T571, V574, S576, S577, L580, G582-K595, W597-L614, A616, F618 |
| WP_249984768.1 | MipA/OmpV family protein | 33 | I13-F15, M17, T18, V22, D26, F27, A30, I34, H40, D48, T52, S79, N80, A94, E99, P102, C143, F162, V184, V187, Q191, T192, S209, N213, D218, V234, N245, L247, T250, S252, R256 |
| WP_108511463.1 | Outer membrane protein transport protein | 37 | L183, A184, L185, A187, A189, Q190, A191, L192, M193, Q194, N195, A196, P197, Q198, L199, A200, A201, Q202, L203, A204, A205, L206, P207, A208, T210, V211, R235, V240, Q257, L260, L262, Q264, V265, S267, T268, Q269, H391 |
| WP_005757734.1 | Outer membrane beta-barrel protein | 18 | F11, I17, D22, S39, N46, Q99, H114, R128, Q133, P149, T157, P159, N162, Q164, Y166, T167, K171, H180 |
| **GROUP B** |  |  |  |
| WP_249984435.1 | TonB-dependent receptor | 455 | M1-V14, M16-V24, A26-F35, S40, E42-S65, Q67, K68, K70-G72, S74, S85, Q90, S95, V99, V100, L108, F115, I118, Q122, L123, I129-V131, T133, I141, A151, N152, P155, I159, Q169, L170, Q172, V177-N180, Y183, N184, N186, H188, K190-V192, A195, L196, F198, G199, G201, K202, I204, V208, E210, L212, G214, H216, H217, H219, L225-K227, L229, N230, V232, T235-S241, N243, Y244, A247, F248, G250-G253, V255, F257, A258, N260, L261, E264, K265, L275, G277, A279, A280, I282-V287, N289-Y291, G294, L295, M300, H301, L305, N307, T308, F310, H311, G313-M318, G320, K321, D325, Y328, D331, I336, A337, L340-K346, Y348, I350, A352-Q356, M358-E362, I364, L366, Y368, E370-R372, K374, D376, D379-I382, V384-F387, N390, L394-F399, A404, I410, A412, Q415-I423, R425-K450, D452-A457, I459, E460, I462, S464, M466, I471, Q473-Q477, L480, L483, V485, T487, E488, D493, E495, R498-W500, F502-V504, R506-A530, F533-R539, T541, S544, A546-S548, N550-T552, D554, T556-S558, I560, Y561, H563, H567, K578, V583, R590, K593, V596, D598, W600, V602, L604, L607, G608, L611, S612, V615-V617, Y619, D621, K623, R625, F627, Q629, T630, H632-S634, L637, S638, N640, N643, S645, K646, Y649, Y650, V652, G654-L660, E663-G667, K676, Y678, P682, T683, R685-Q701, P707, M709, F713, V715-M719, S722, T724-E728, Y733, Q734, K736, V738, P740, L741, N743-S749, L755, D756, S758, H759, A762-N765, Q767-A772, V775, R778-I780, S782, F786, F789, V797, T798, G800-H804 |
| WP_249984789.1 | TonB-dependent receptor | 116 | S17, Q18, T30, P64, V168, E176, A203, L245, A286, Q340, N353, A378, A402, V456, V460, V462, E468, L473, E517, R518, V523, F525, P532, A551, K557, D569, V594, D597, T600, T602, K606, G607, N630, A638, K641, Y646, A649, I650, N661, A664, E670-V672, D674-I680, K683, M705, E712, L717, S719, K726, T728, R738-S748, E750-F797 |
| WP_324023022.1 | TonB-dependent receptor domain-containing protein | 33 | M1, Q28, A126, I157, K237, A339, K400, K427, V429, A430, T488, N505, E532, L541, A592, F640, L662, S698, C717, A737, D771, K782, D789, A790, R792, E793, D796, G805, A809, V831, I833, V839, L879 |
| WP_005756417.1 | Toxin/drug exporter Tde | 18 | S14, Q34, M39, H42, K100, A101, R116, G120, V178, A212, A248, K252, G288, A289, S309, A315, N359, A363 |
| WP_104227660.1 | Efflux transporter outer membrane subunit | 13 | V8, G16, T43, H71, G98, Q193, T288, Y294, T402, D409, Q438, T447, V458 |
| WP_083005347.1 | OmpW family outer membrane protein | 4 | F6, G46, L132, L180 |
| WP_249984615.1 | Sialidase family protein | 648 | P4, V5, L7, L8, L10, I13, T15, M17-G44, T46, N48, G51, P53-G64, T66-L69, Y71, S72, A76-I79, D81, N83-F89, L91, N94-S97, Q99-R101, P104-A107, E110, G112, H114, S115, K117, K118, A121-D126, K128-M135, P137-D145, S147-Y169, N171-K181, T183, G185, L187, S188, Q190-R201, I203-G217, L219, G220, V222, T224, V227-D230, T232, V234, Q238-D243, I245-Y251, K253, N255, S258, M261-F280, I282, D283, V287, T289-E309, L311, K313, H316-P320, N322-I337, V339-S343, R346-V350, K352-N354, N356, N358, G360-R363, N365, A367, L368, M370-A383, I385-A393, A395, G396, L400, Y402-E404, N406-I409, F411, D413-G415, I417, V419-A428, E431-E435, G437-I442, T444-V446, N450-K457, Q459, K460, A462, S464-V480, N482, R483, M485-E491, T493-E496, K498, S499, A501-L506, S508-L535, I537-T553, L555, F557-K559, I561-Q563, K566, N568-I574, Y576-V582, E585, D587-T598, A600-F603, T605-A613, V615-H619, N621-Q624, V626-F629, V631-D639, V641-F645, R647-V651, D654-V656, D658-A670, V673, N674, D676, H678-A687, T689, S693-V703, K705-V713, L715, I717, R718, L721-S725, S727-S742, A744-Q747, N749, Q750, I753-A761, N763-F770, H773, L774, V776-L782, K784-V792, F794-S799, H801, W802 |

≠ Proteins were named as per their annotation in the NCBI database, searched using protein accession number.

**Table S5**: Top 20 conformational B-cell epitopes selected for the identified proteins.

| **Protein accession no.** | **Residues** | **Number of residues** | **Score** |
| --- | --- | --- | --- |
| WP_104228050.1 | A:E2, A:S3, A:P4, A:K5 | 4 | 0.996 |
| WP_126417754.1 | A:V2, A:Y3, A:P4, A:M5 | 4 | 0.996 |
| WP_249984729.1 | A:M1, A:K2, A:L3, A:K4 | 4 | 0.996 |
| WP_096743251.1 | A:S2, A:F3, A:K4 | 3 | 0.993 |
| WP_249984435.1 | A:M1, A:I2, A:S3, A:R4 | 4 | 0.993 |
| WP_324023075.1 | A:S4, A:S5, A:L6 | 3 | 0.992 |
| WP_324023022.1 | A:M1, A:V2, A:T3, A:N4, A:K5, A:L6, A:I7, A:K8, A:Q9, A:I10, A:T11, A:Y12, A:L13, A:L14, A:P15 | 15 | 0.991 |
| WP_126417535.1 | A:Y3, A:P4, A:L5, A:S6, A:Y7, A:K8, A:N9, A:I10, A:A11, A:R12 | 10 | 0.99 |
| WP_249984830.1 | A:C4, A:L5, A:I6, A:I7, A:Y8 | 5 | 0.989 |
| WP_324023075.1 | A:F7, A:V8, A:F9 | 3 | 0.987 |
| WP_096743251.1 | A:K6, A:T7, A:L8, A:A9, A:L10, A:F11 | 6 | 0.986 |
| WP_249984830.1 | A:L9, A:N10, A:S11, A:I12, A:L13, A:I14, A:C15, A:L16, A:F17, A:F18, A:S19, A:F20, A:Q21, A:T22, A:W23 | 15 | 0.986 |
| WP_005756417.1 | A:N5, A:P6, A:L7, A:T8, A:F9 | 5 | 0.984 |
| WP_126417535.1 | A:S13, A:I14, A:P15, A:F16 | 4 | 0.981 |
| WP_249984417.1 | A:M1, A:M2, A:N3, A:L4, A:K5, A:P6, A:L7, A:T8, A:F9 | 9 | 0.981 |
| WP_249984768.1 | A:N3, A:S5, A:K6 | 3 | 0.978 |
| WP_249984417.1 | A:L19, A:S20, A:G21, A:V22, A:I23, A:Y24 | 6 | 0.976 |
| WP_324022762.1 | A:K3, A:A4, A:K5, A:G6, A:I7, A:N8, A:L9, A:Y10 | 8 | 0.973 |
| WP_249984417.1 | A:L16, A:V17, A:F18 | 3 | 0.971 |
| WP_324022736.1 | A:M1, A:K2, A:L3, A:L4, A:S5, A:N6, A:T7, A:K8, A:A9, A:S10, A:L11, A:I12, A:T13, A:K14, A:T15, A:V16, A:F17 | 17 | 0.968 |

**Table S6:** Comparative binding energy analysis of MEV construct with OmpH (positive control).

| **Complex** | **MEV construct** | | **OmpH (positive control)** | |
| --- | --- | --- | --- | --- |
|  | **ΔG (kcal/mol)** | **K_d_ (M)** | **ΔG (kcal/mol)** | **K_d_ (M)** |
| TLR4 and MD2 coreceptor | -23.3 | 7.7 × 10^-18^ | -19.9 | 2.4 × 10^-15^ |
| TLR2 | -20.7 | 6.3 × 10^-16^ | -18.0 | 6.1 × 10^-14^ |

**Supplemental Figures**

MGIINTLQKYYCRVRGGRCAVLSCLPKEEQIGKCSTRGRKCCRRKKEAAAKAKFVAAWTLKAAAAAYFVIDNYLSKAAYYQYDTRLNKAAYSTHYRVPAFAAYYLKHYPFLMAAYMKYNDSWTFGPGPGGLNAVHAKAKLERYAGPGPGDSRLTYSSKLIRGESGPGPGKENSTYGRSSSWSLAGPGPGPSYRAEMASYANNLYGPGPGEIAYTHTDSRLTYSSKKNLTSPMPGFAYKNKKNHTLEYRKKKHHHHHH

**Fig. S1 Amino acid sequence of the MEV construct.**

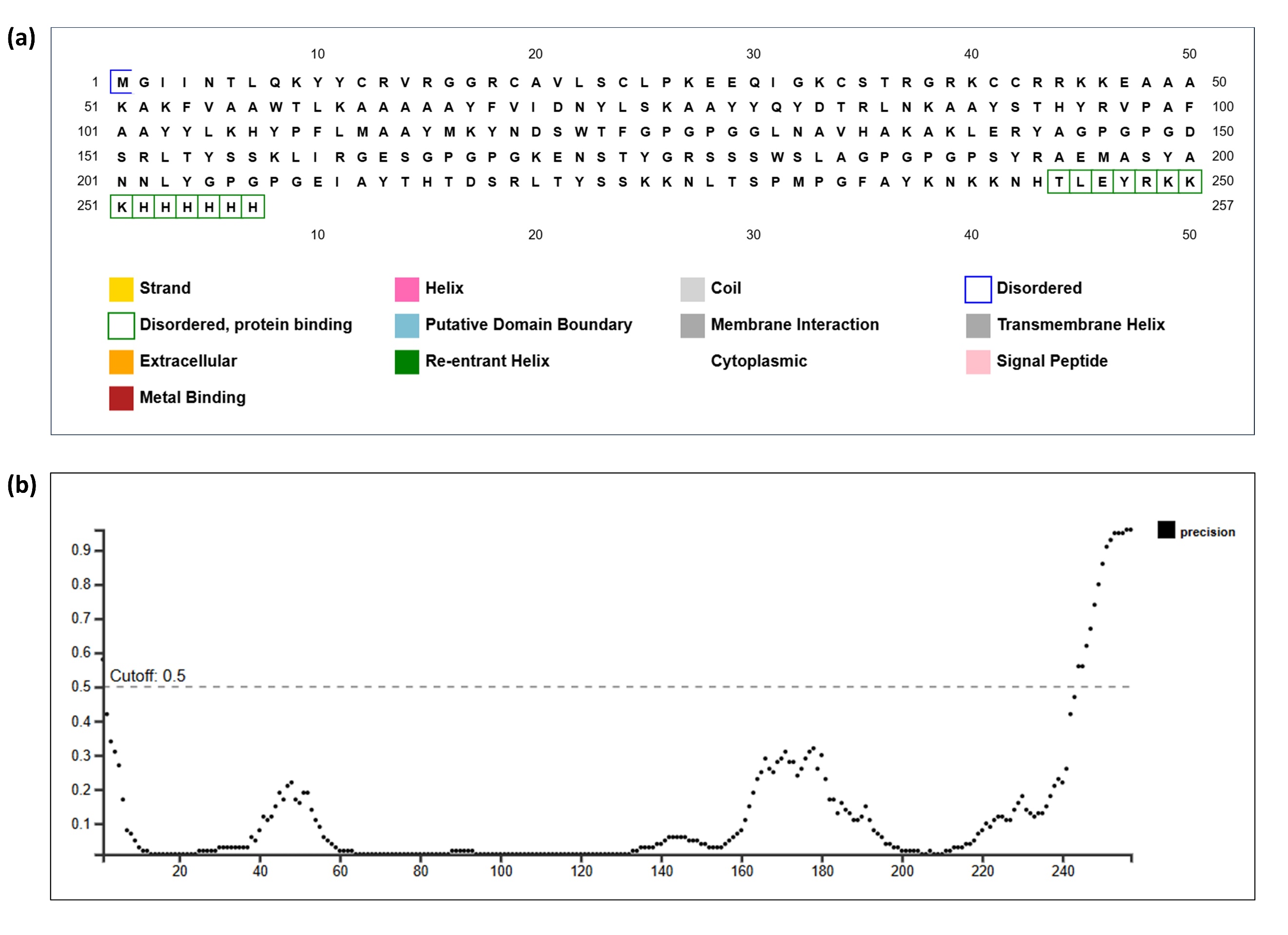
**Fig. S2 Disordered region prediction of the MEV construct.** **(a)** Predicted disordered regions in the MEV. **(b)** Residue-wise disorder propensity profile with a prediction cutoff of 0.5 indicated by the dashed line.

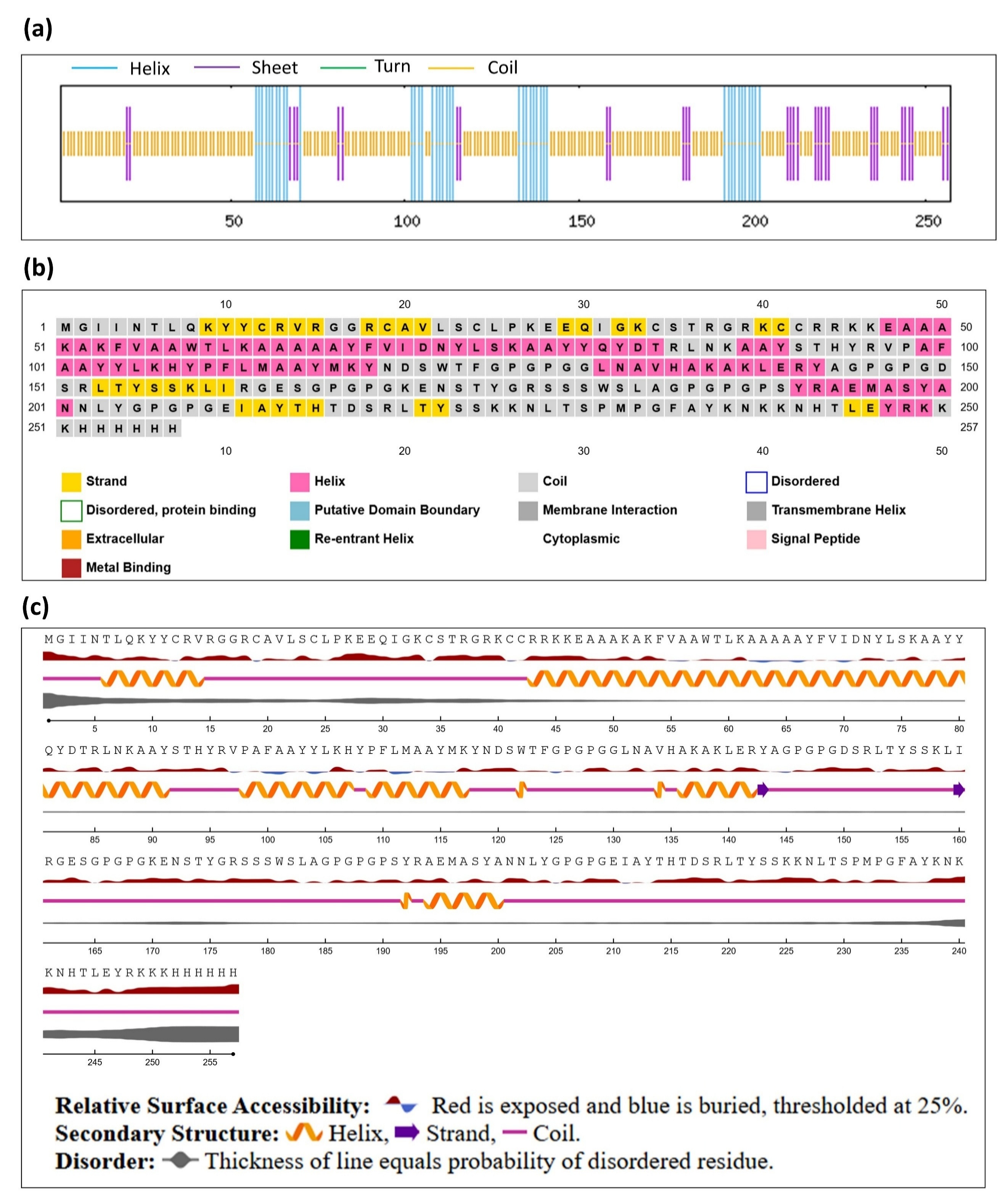

**Fig. S3 Secondary structure prediction of the MEV using (a)** SOPMA, **(b)** PSIPRED, and **(c)** NetSurP

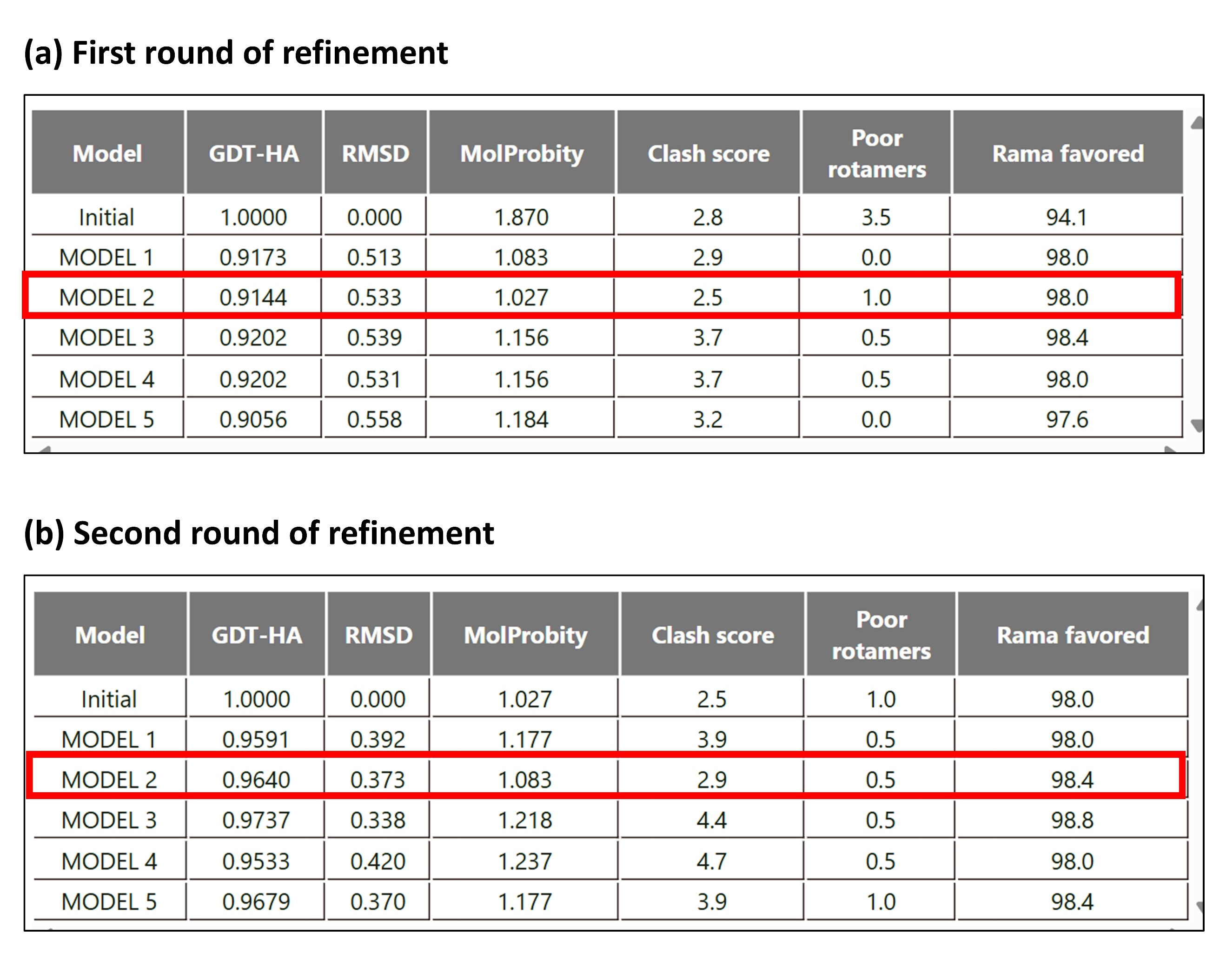
**Fig. S4 Structural refinement of the MEV model using GalaxyRefine. (a)** First round of refinement, with Model 2 (red box) chosen for subsequent refinement. **(b)** Second round of refinement, with Model 2 (red box) selected as the final refined model based on its overall structural quality.

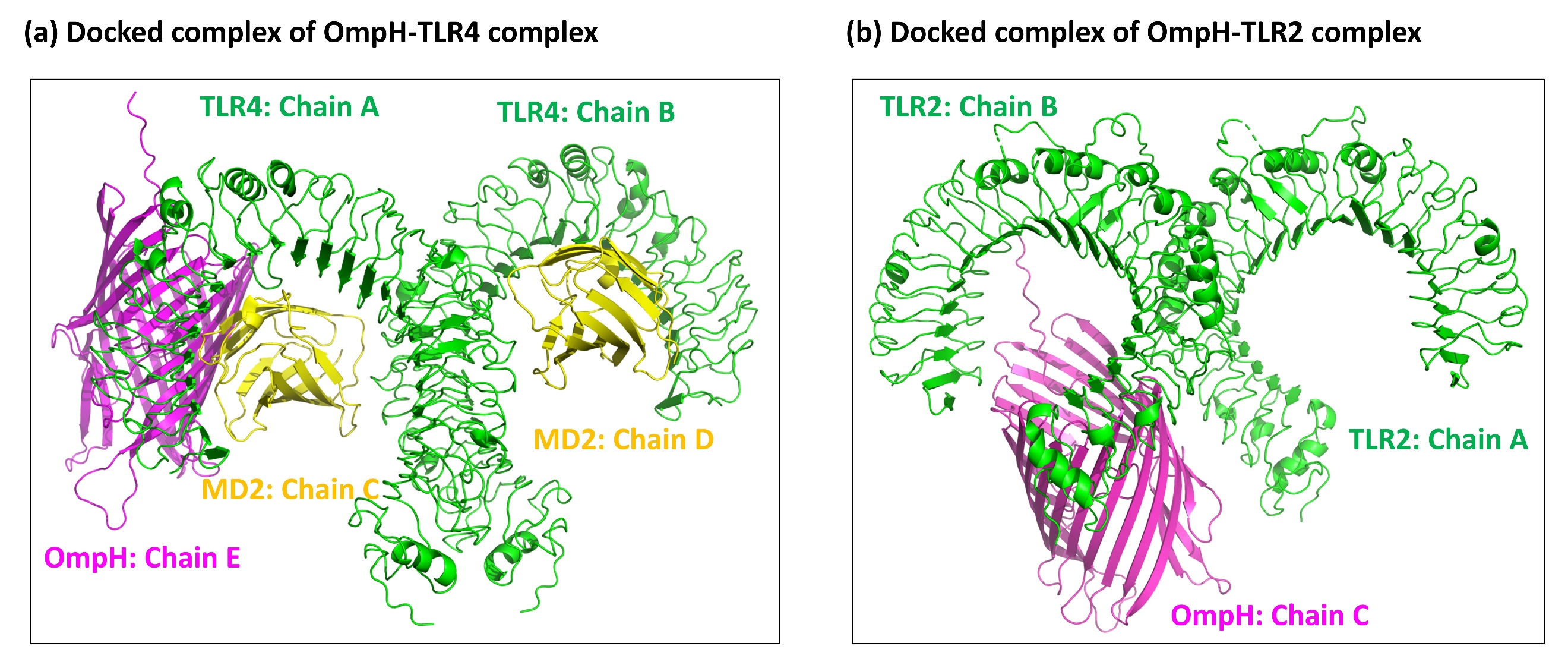

**Fig. S5 Molecular docking of OmpH with TLR4 and TLR2. (a)** The docked complex of OmpH with TLR4 (PDB id: 3FXI), along with the co-receptor myeloid differentiation factor 2 (MD2). **(b)** Docked complex of the OmpH with TLR2 (PDB id: 6NIG).

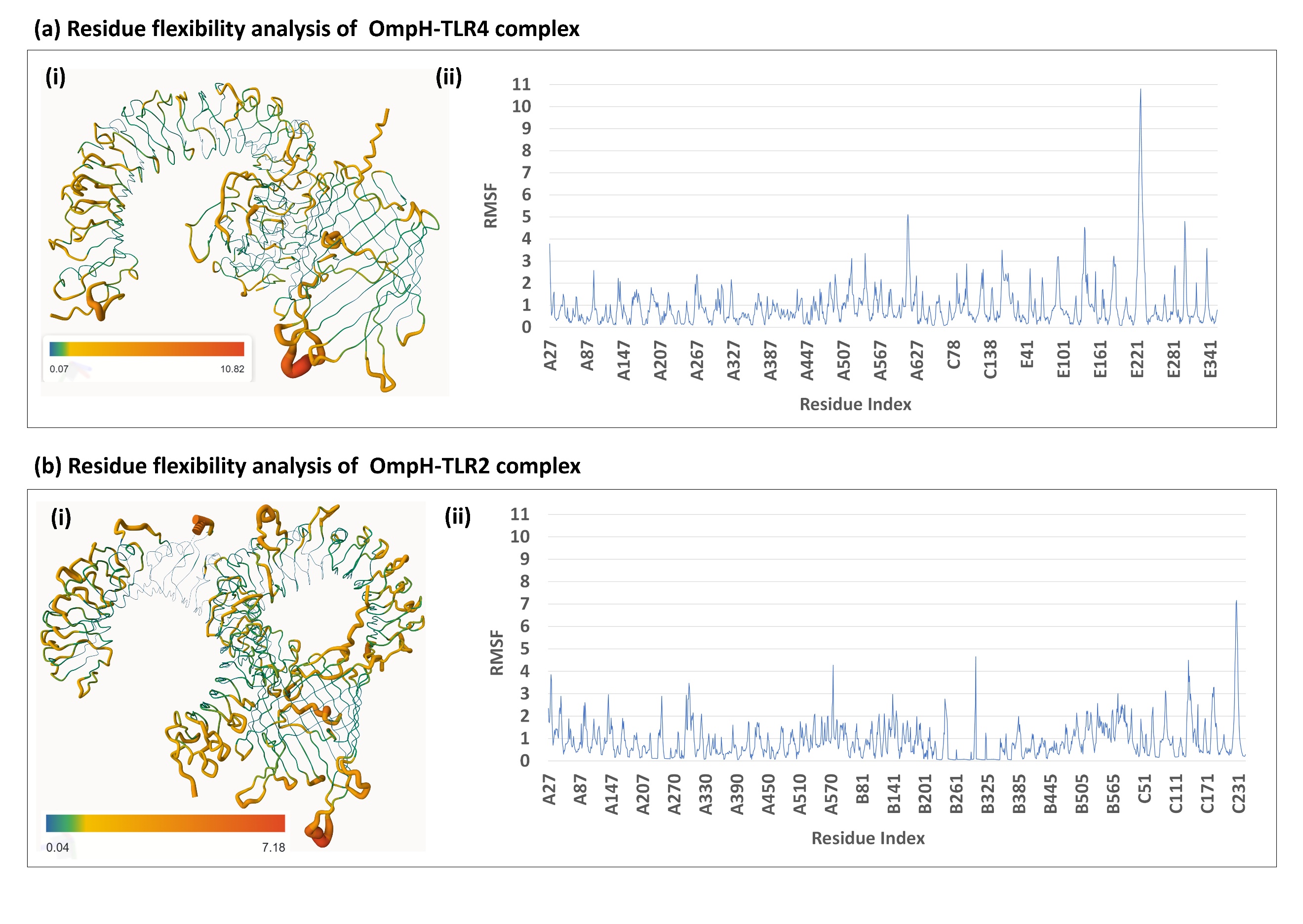

**Fig. S6 Residue flexibility analysis of OmpH-TLR4 and OmpH-TLR2 complexes**. **(a)** OmpH-TLR4 complex and **(b)** OmpH-TLR2 complex. **(i)** Structural representation of residue flexibility in the protein-ligand complex. **(ii)** RMSF profiles of the corresponding complexes throughout the simulation. Blue and red regions indicate stable residues and highly flexible residues, respectively.

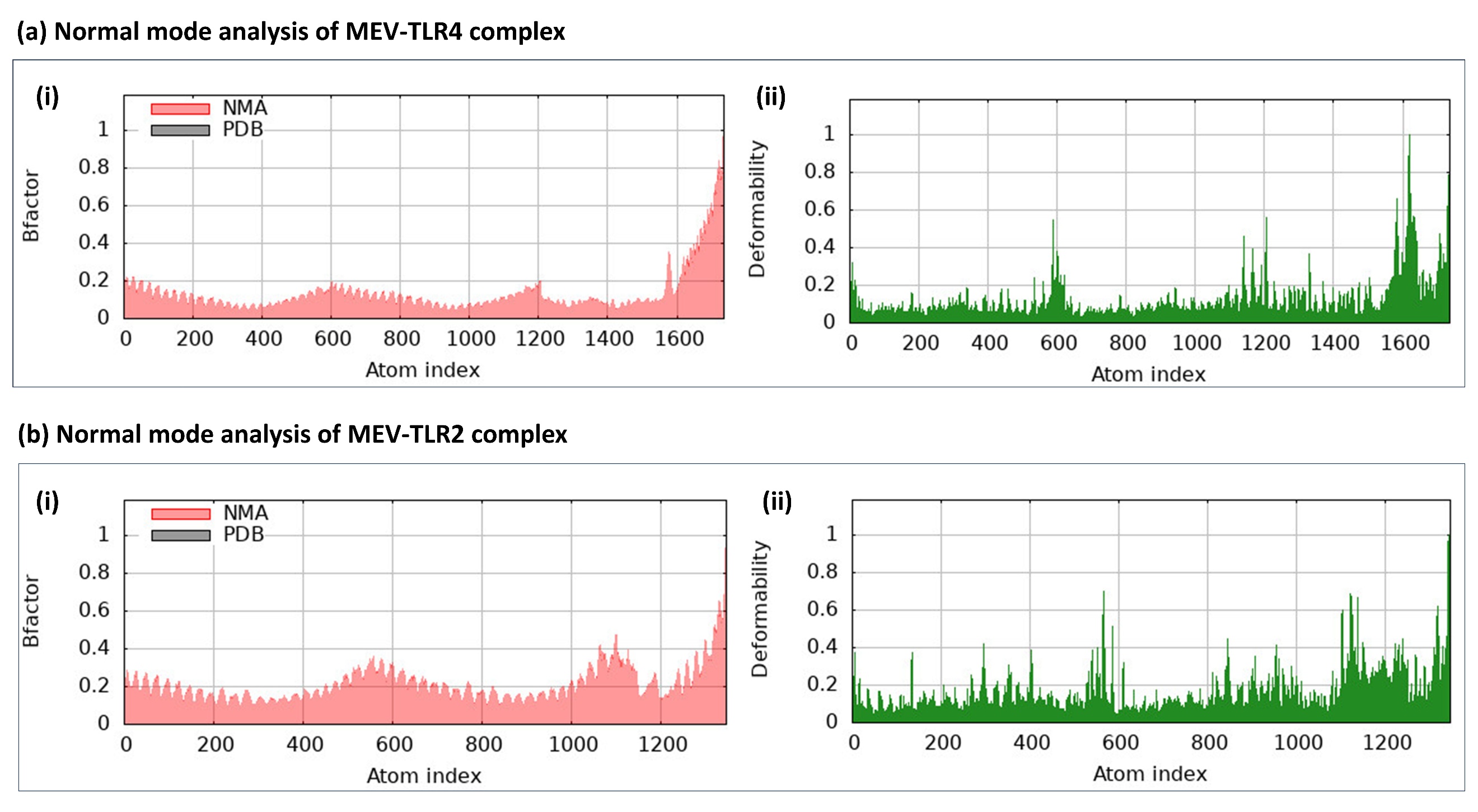

**Fig. S7 Normal mode analysis.** **(a)** MEV-TLR4 complex and **(b)** MEV-TLR2 complex. **(i)** B-factor comparison **(ii)** Deformability plot

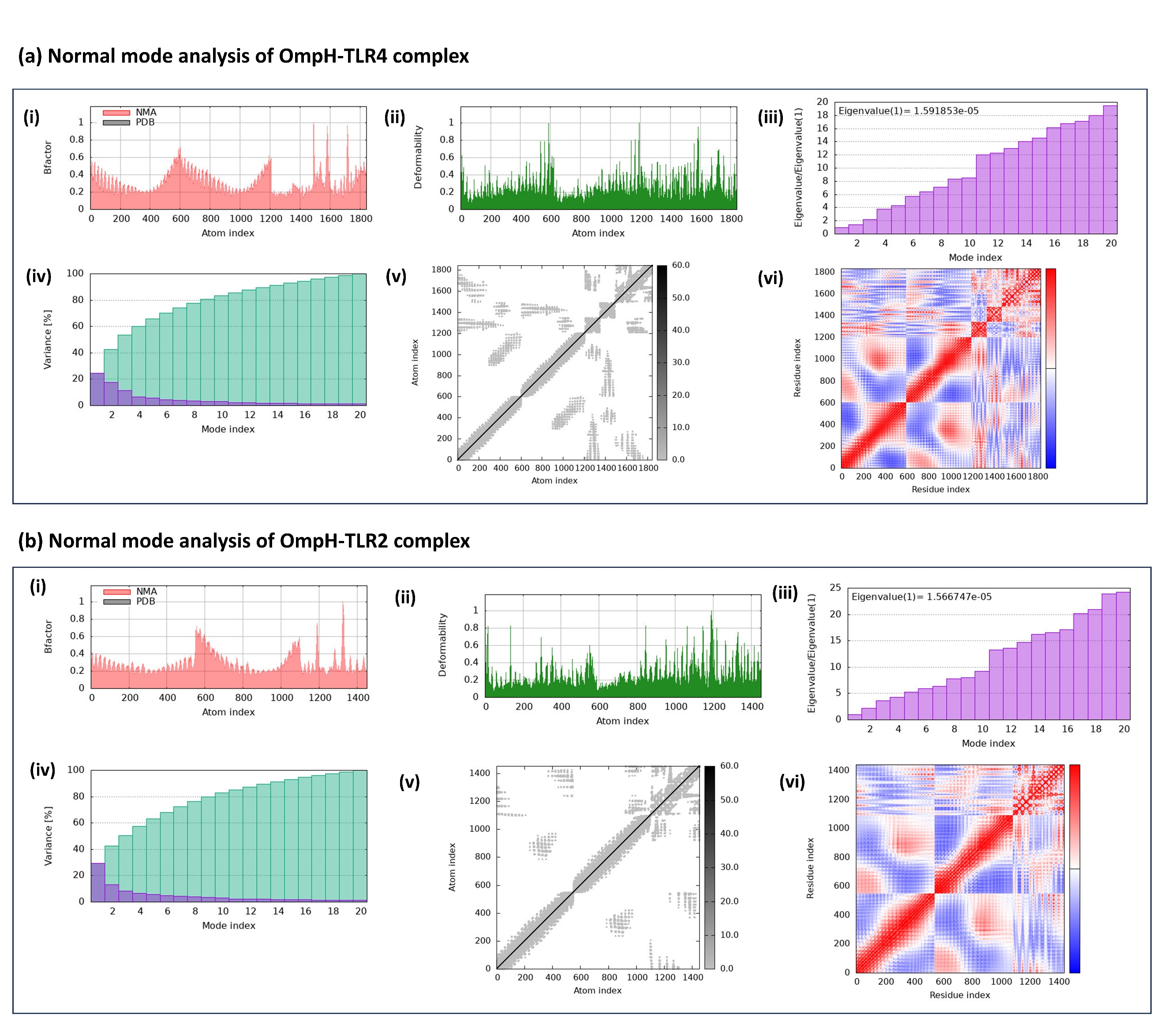

**Fig. S8 Normal mode analysis of OmpH-TLR4 and OmpH-TLR2 complexes. (a)** OmpH-TLR4 complex and **(b)** OmpH-TLR2 complex. **(i)** B-factor comparison **(ii)** Deformability plot **(iii)** Eigenvalue spectra **(iv)** Variance plot **(v)** Elastic network model **(vi)** Covariance map

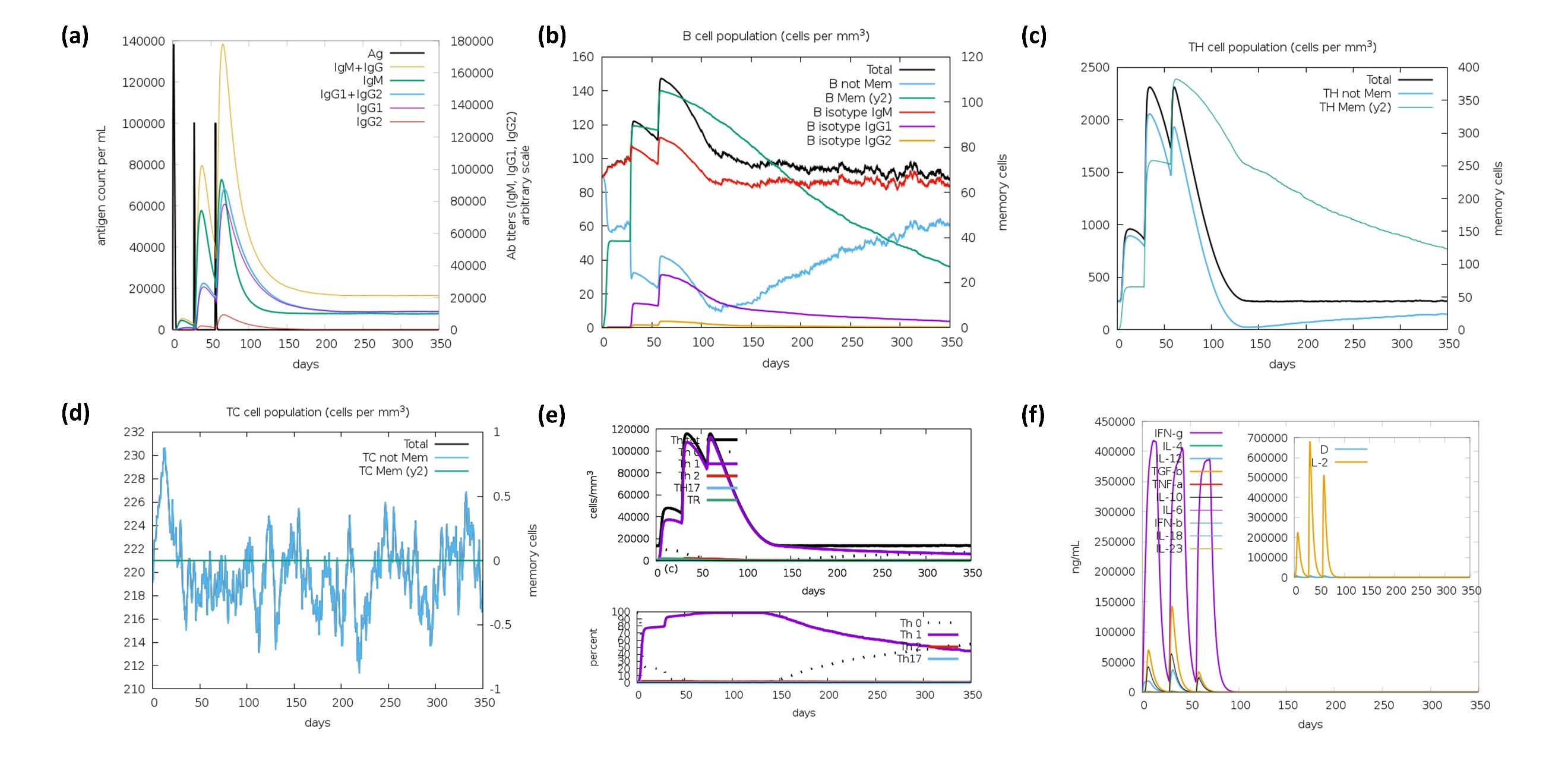
**Fig. S9 In silico immune simulation of OmpH.** **(a)** Antigen and subtypes of immunoglobulin levels. **(b)** B-cell isotypes in various states. **(c)** Helper T-cell population **(d)** Cytotoxic T-cell population **(e)** Helper T-cell isotypes in various states. **(f)** Concentration of cytokines and interleukins.
